# Object Detection Networks and Augmented Reality for Cellular Detection in Fluorescence Microscopy Acquisition and Analysis

**DOI:** 10.1101/544833

**Authors:** D Waithe, JM Brown, K Reglinski, I Diez-Sevilla, D Roberts, Christian Eggeling

## Abstract

In this paper we demonstrate the application of object detection networks for the classification and localization of cells in fluorescence microscopy. We benchmark two leading object detection algorithms across multiple challenging 2-D microscopy datasets as well as develop and demonstrate an algorithm which can localize and image cells in 3-D, in real-time. Furthermore, we exploit the fast processing of these algorithms and develop a simple and effective Augmented Reality (AR) system for fluorescence microscopy systems. Object detection networks are well-known high performance networks famously applied to the task of identifying and localizing objects in photography images. Here we show their application and efficiency for localizing cells in fluorescence microscopy images. Object detection algorithms are typically trained on many thousands of images, which can be prohibitive within the biological sciences due to the cost of imaging and annotating large amounts of data. Through taking different cell types and assays as an example, we show that with some careful considerations it is possible to achieve very high performance with datasets with as few as 26 images present. Using our approach, it is possible for relatively non-skilled users to automate detection of cell classes with a variety of appearances and enable new avenues for automation of conventionally manual fluorescence microscopy acquisition pipelines.

## Introduction

Even the most advanced light microscopes typically have a wide-field modality that visualises a sample specimen and allows the user to view it by looking down the eyepiece. The user will look down the binocular, find an area to image and then commit the system to acquiring an image within that region. Subsequent to this, the way in which the measurement takes place is highly flexible, it maybe a sequence of images to form a time-series, a volumetric stack, a super-resolution microscopy image or some kind of spectroscopic readout. The process of identifying cells for imaging is challenging in that it requires domain specific knowledge, but is also repetitive, in that the process involves localising many cells through looking down the eyepiece. This is time-consuming for the scientists who must perform the acquisition and who must be present throughout much of the experiment.

There are a number of high-content automated optical light microscopes which can find and image cells on the fly, relieving the effort of acquisition for the researchers and providing thorough documentation of the acquisition pipeline [1, 2]. Typically, these systems don’t intelligently identify cells before imaging, but rather, exhaustively image a plate or slide and then perform analysis on all the samples imaged. This dense sampling is perfect for obtaining images of all the cells present, but, due to the assay format and the hardware required, means that subsequent specialised imaging is hard to perform. These systems are ideally suited to relatively static assays, with a high level of repetition, and relatively low magnification. A system like this for example would test a large library of compounds acting on cultured cells and the activity would be established through visualisation of a reporter or through morphological analysis of the cells being imaged [1, 2]. Techniques like this are often built on signal-processing methods and are utilised for two-dimensional (2-D), or more recently, three-dimensional (3-D) imaging and analysis of cells and tissues [3]. Once created, these methods are powerful and fast, but they lack flexibility; they cannot be easily tuned to different cell types or assays without skilled input. Through basic optimisation these algorithms can be applied to conventional microscopes that have been retrofitted with an automated stage. The problem however is the fragility of these algorithms when applied to a new setting (e.g. a different staining or microscope). A skilled analyst is often required to tweak the parameters or to modify the algorithm to detect a different cellular appearance. An optimum solution would be an algorithm that is easy to adapt by a relatively unskilled user to recognise and localise cells of any type, reproducibly and reliably and across different microscopes.

One of the key benefits of manual microscopy is, as mentioned, the specificity through it can be applied and the potential for customisation. One of the pitfalls however with a highly manual approach is that decisions made through the experimental pipeline are difficult to describe, illustrate and therefore to document and share. This lack of ability to communicate decisions made during the acquisition process means it is difficult for the scientific community as a whole to question and discuss methodologies and selection strategies and which means that we are at risk as a community from unconscious (and potentially conscious) bias of scientists. This issue is difficult to address but can be approached by adopting and introducing technology that allows improved documentation and reproducibility of data acquisition during an experiment. This is a delicate balance however as it must be done without reducing the flexibility of the manual microscopy on which the automation is applied or restricting the workflow or experiments performed by the scientist. This can be achieved, as we will see, through introducing more automation into the experimental pipeline and through using advanced cellular detection methodologies.

Computer vision (CV) has developed to solve various challenges in video and photography. From this field, high-performance algorithms have evolved which perform classification and localization in images even when there is a high degree of variability and complexity in object appearance. Recent advances in machine learning and computer hardware have made possible a new generation of end-to-end neural networks with unparalleled performance [4]. In recent years algorithms inspired from the CV domain have made a noticeable impact in the domain of microscopy image analysis, and interest continues to grow [5–8]. Object detection, a sub-discipline of CV, has developed with the goal of predicting bounding boxes for multiple objects in images or videos for potentially multiple classes at different scales. In the past, this approach has been applied across many fields including pedestrian detection, face detection, autonomous vehicles, and for robotics (for reviews: [9–15]). So far object detection algorithms have not been used extensively for microscopy based applications, though there are some recent contributions which utilize these type of networks (e.g. for Astrocyte detection) [8, 16]. In this article we compare two state-of-the art object detection frameworks from the CV domain, Faster-RCNN and YOLO, and evaluate their performance for identifying and localizing cells in microscopy images, and show how they can be used to explore a 3-D space rather than being applied only to static 2-D images.

Faster-RCNN (Region-based Convolutional Neural Network) was the first network to combine features for region proposal with object classification and represents the culmination of a systematic set of advances and optimizations [17],[18],[19]. This network, and subsequent networks, leverage the depth and power of neuronal networks to learn a transformation between the input space (the image) and the output space, in this case, a set of bounding boxes for the objects located within the input image. All object detection networks employ layers that calculate features across an image. These feature-calculating layers of the network are structurally identical to those used in classification networks, and they are generally composed of multiple successive convolution, activation and pooling layers whose parameters (or weights) dictate how the features are calculated. Once trained and applied to an image, the features numerically describe the shape and form of the objects and shapes present in an image. From this feature representation, multiple ‘anchor’ regions are spawned that are then evaluated in their capacity to bound objects in the input space. Through optimizing the size and placement of these regions, with respect to the objects in the input training images it is possible to identify and localize objects using the information encoded in the feature representation. The network weights are trained to optimize this process for finding and ultimately classifying the objects within the regions of the image. Once classified, the bounding boxes are further modulated, through class specific layers that further refine the bounding box positions for each object. Following Faster-RCNN, several other competitive algorithms have been developed which compete with and out-perform Faster-RCNN in several aspects (e.g. SSD [20], YOLO[21–23]).

YOLO (You Only Look Once) is a competing framework for object classification, which improves on Faster-RCNN in terms of accuracy and speed. For this study YOLOv2 and YOLOv3 have been benchmarked [23]. The YOLO framework has consistently been found to be the fastest network for object detection compared to its competitors like Faster-RCNN and SSD, and it is also highly accurate [23]. Like Faster-RCNN, YOLOv2 utilizes convolutional anchor boxes as the basis of the object detection, but has a different overall structure that yields superior performance. Amongst the differences which are summarized in [21], YOLO employs a technique known as batch normalization, at each layer, which is a recent but proven method for regularization, shown to improve the accuracy of many networks [24]. In this study, we thoroughly characterize YOLOv2 algorithm and compare it to Faster-RCNN to establish the performance of both algorithms on relatively small datasets in this different modality. We additionally benchmark YOLOv3 but we find this network does not perform as well as the other networks on this data.

As mentioned, object detection algorithms are derived from work performed on photography images. There are several qualities of microscopy images, specifically in fluorescence microscopy, that make them distinct from photography images, and that could be important in terms of the performance of the proposed algorithms. Firstly, occlusion is minimal; for example, signal in fluorescence microscopy is additive and it is usually possible to make out structures even when cells overlap, unlike in photography images where occlusions will hide objects. Secondly, there is no perspective or scale issues as the depth-of-view in microscopy is typically narrow. There are however challenging aspects of microscopy images; cells will have a variable intensity, for example, some cells may be bright and others darker within the same field-of-view. Furthermore, the appearance can vary dramatically between cells due to natural variation in morphology and shape. Additionally, cells may be densely packed or even overlap, making it complicated to delineate individual cells. This makes the question of whether object detection networks are effective or not for microscopy an important one. What potentially makes the object detection networks so attractive to microscopy is their accuracy, their ease of use, and their predictive speed. Thanks in part to the design of the more recent algorithms, these algorithms can be easily implemented efficiently on a GPU and so can evaluate images in real-time. This makes them perfect for use in an automated microscopy setup where a microscope will image and apply analysis in sequence. Annotating and manually segmenting cells for example fluorescence images can be very demanding and can often require pixel level segmentations to be drawn. Object detection algorithms on the other-hand utilize bounding boxes, which are relatively simple for a user to provide through a simple and fast annotation procedure. For these reasons, the possibility of using object detection algorithms in microscopy is an interesting one and worthy of investigation.

In this paper we show that Faster-RCNN and YOLOv2.0 behave very well as object detection algorithms for fluorescence microscopy. We show that despite their complexity these algorithms can be trained to work on relatively modest sized training datasets. Furthermore, we prototype and test an algorithm that can utilize the bounding boxes predicted by these algorithms to find and localize cells in a 3-D environment, a framework we call the Autonomous Microscope Control Algorithm (AMCA).

## Materials and Methods

### Dataset generation

To create a collection of datasets to test the capabilities of the object detection algorithms a diverse range of cell types were assembled and stained with commonly used dyes. The data in its entirety as well as the annotations are available in the repository (https://doi.org/10.5281/zenodo.2548493).

### Erythroblast DAPI (+glycophorin A)

erythroblast cells were stained with DAPI and for glycophorin A protein. Cells were stained with CD235a antibody (JC159 clone from Dako) and with Alexa Fluor 488 secondary antibody (Invitrogen). DAPI staining was performed through using VectaShield Hard Set mounting solution with DAPI (Vector Lab). Num. of images used for training: 80 and testing: 80. Average number of cells per image: 4.5.

### Neuroblastoma phalloidin (+DAPI)

images of neuroblastoma cells (N1E115) stained with phalloidin and DAPI were acquired from the Cell Image Library [25]. The images were stained for FITC-phalloidin and DAPI. Num. of images used for training: 180, testing: 180. Average number of cells per image: 11.7.

### Fibroblast nucleopore

fibroblast (GM5756T) cells were stained for a nucleopore protein. Staining was performed using Anti-Nup153 mouse antibody (Abcam) and detected with anti-mouse Alexa Fluor 488. Num. of images for training: 26 and testing: 20. Average number of cells per image: 4.8.

### Eukaryote DAPI

eukaryote cells were stained with DAPI and fixed and mounted in Vectashield. Num. of images for training: 40 and testing: 40. Average number of cells per image: 8.9.

### C127 DAPI

C127 cells were initially treated with a technique called RASER-FISH[26], stained with DAPI and fixed and mounted in Vectashield (Vector Lab). Num. of images for training: 30 and testing: 30. Average number of cells per image: 7.1.

### HEK peroxisome

HEK-293 cells expressing peroxisome localized GFP-SCP2 protein. Cells were transfected with GFP-SCP2 protein, which contains the PTS-1 localization signal, which redirects the fluorescently tagged protein to the actively importing peroxisomes[27]. Cells were fixed and mounted. Num. of images for training: 55 and testing: 55. Average number of cells per image: 7.9.

### Dataset Annotation

Datasets were annotated by a skilled user. These annotations represent the ground-truth of each image with bounding boxes (regions) drawn around each cell present within the staining. The dataset labels were then converted into a format compatible both with Faster-RCNN (Pascal) and with YOLOv2 and YOLOv3. The scripts used to achieve this are located in the repository (https://doi.org/10.5281/zenodo.2594644).

### Microscopy setup

With the exception of the neuroblastoma and erythroblast cell datasets all images, were acquired on an Olympus IX73 microscope with a 100X UPlanSApo, NA 1.4 objective. The microscope was also equipped with a Photometrics Prime sCMOS camera, a CoolLED Ultra pe300 LED light source, an ASI automated XY stage and a PI Piezo (P-733 2CL). The erythroblast dataset was acquired on a DeltaVision Elite (GE Healthcare Life Sciences) equipped with an Olympus 60x NA 1.42 lens, filters for DAPI (exc. 390 nm, emi. 435 nm) and FITC (exc. 475 nm, emi 525 nm) and a CoolSNAP HQ2 camera. The neuroblastoma phalloidin +DAPI cell line was acquired on a Zeiss Aviovert 200 microscope with filters for DAPI and FITC [25].

### Augmented Reality modifications

To develop the augmented reality setup we adapted commercial components with custom parts. The augmented reality effect is created through the merging of the image emanating from a screen projecting graphics with the image emanating from the microscopy sample. This was achieved through the coupling of a 50:50 beamsplitter (Thorlabs, BSW10R) into the light path of the microscope. This was realized through adapting a Mightex Dichroic/filter cube (DSI-CUBE-OL-UA) to fit in between the observation tube and the observation tube mount of an IX73 microscope (Olympus). The Mightex Dichroic filter has the required circular dovetail mounts to fit within the binocular of the IX73 system, but in its default configuration the beamsplitter couples light toward the specimen and not the observer, which is what we require for the augmented reality system. To correct this we engineered two adapter plates to reverse the gender of the mounts, details of these plates can be found in the supplementary materials (SM1). The computer screen (Pimoroni, HDMI 8” IPS LCD Screen Kit) was positioned to the right of the microscope so that the base of the screen was parallel to the Mightex cube and perpendicular to the light path through the microscope. The screen was secured at the desired angle using standard M6 Post components (Thorlabs) and an Ailun Tripod Mount Adapter (Amazon, B071XHYG5R). Attached to the Mightex cube in between the beam splitter and light coming from the computer screen was a 300mm biconvex lens (Thorlabs, LB1779) which converged the light from the screen onto the beam splitter. The computer screen was placed around 30cm from the screen which resulted in an in focus view of the screen graphics when looking down the binocular. An optional modification we made to the conventional IX73 setup was to use a 50:50 beamsplitter cube in place of the mirror that directs light either to the camera or to the binocular. We made this modification so we could simultaneously view the specimen down the binocular and also record the same image on the computer. For this we engineered our own cube holder (SM2) and using superglue adhesive attached a 30mm 50R/50T Standard Cube Beamsplitter (#32-701, Edmund Optics Ltd).

### Computer Hardware

Benchmark experiments were run on Dell PowerEdge R730 Server: 2x Intel Xeon E5-2650, 256 RDIMM RAM, NVIDIA Tesla K80 GPU with CentOS 7 installed. Real-time acquisition experiments were run on Dell Precision Tower, with 32GB RDIMM Ram, Nvidia Quadro P5000 16GB GPU, Dual Xeon Processor E5-2637 with Windows 10 installed.

### Object Detection Algorithms

In this study we took two leading publically available object detection network frameworks (Faster-RCNN and YOLO). Because of the way in which these different frameworks function we had to ensure that the datasets were correctly formatted in each case. Each dataset has annotations in form of the PASCAL dataset [28], which is converted using scripts into a form compatible with YOLO. Each network is presented with one or more datasets for training.

The code used for the Faster-RCNN is a tensorflow implementation and was modified from https://github.com/dBeker/Faster-RCNN-TensorFlow-Python3.5 and can be found here https://zenodo.org/record/2594642. Faster-RCNN was configured as follows. The VGG16 network was used to initialize the classification layers. The parameters for learning were configured as follows: ‘Weight_decay’ = 0.0005, ‘learning_rate’ = 0.001, ‘momentum’ = 0.8, ‘gamma’ = 0.1, ‘batch_size’ = 256, ‘max_iters’ = 40000, ‘step_size’ = 30000. The network was modified to flip images not just horizontally but vertically during data augmentation.

YOLOv2 was cloned from the source (https://pjreddie.com/darknet/yolov2/) and was modified for this work (https://doi.org/10.5281/zenodo.2594648). The modified YOLOv2 network was configured with the following settings: ‘batch’ = 64, ‘subdivisions’ = 8, ‘height’ = 416, ‘width’ = 416, ‘channels’ = 3, ‘momentum’ = 0.9, ‘decay’ = 0.0005, ‘angle’ = 0, ‘saturation’ = 1.5, ‘exposure’ = 1.5, ‘hue’ = .1, ‘learning_rate’ = 0.001, ‘burn_in’ = 1000, ‘max_batches’ = 10000, ‘policy’ = steps, ‘steps’ = 4500, 4800, ‘scales’ = .1, .1. The network was modified to flip images not just horizontally but vertically during data augmentation.

YOLOv3 was cloned from the source (https://github.com/pjreddie/darknet) and run with settings: ‘batch’=64, ‘subdivisions’=16, ‘width’=608, ‘height’=608, ‘channels’=3, ‘momentum’=0.9, ‘decay’=0.0005, ‘angle’=0, ‘saturation’ = 1.5, ‘exposure’ = 1.5, ‘hue’=.1, ‘learning_rate’=0.001, ‘burn_in’=1000, ‘max_batches’ = 20000, ‘policy’=steps, ‘steps’=4500,4800, ‘scales’=.1,.1. The network was modified to flip images not just horizontally but vertically during data augmentation. The number of classes was set to 6 and the filters adjusted accordingly to 33 (filters=(classes + 5)*3).

Average Precision (AP) is a commonly used metric for assessing the accuracy of algorithms that are performing classification and/or localization. For this study we use the updated VOC2010 Average Precision definition described in [28] and as follows: In a given 2-D image, containing one or more objects (i.e. cells) a trained object detection network will predict bounding regions for each of the objects contained within the image and associate a level of confidence (0-1.0) with that prediction. At a low confidence threshold many regions will be predicted whereas a higher confidence much fewer, normally for visualization of results we show the predictions with a specific cut-off for each algorithm. For comparison between algorithms however we need to evaluate performance across confidence levels. The first stage in this process is to assess which of the predicted regions (Bp) is overlapping the ground-truth regions (Bgt). Those detections which have an overlap coefficient (ao) of more than 50% are considered correct detections (True Positives, TP) otherwise they are defined as False Positives (FP).

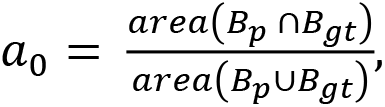

where *B*_*p*_ ∩ *B*_*gt*_ is the intersect and *B*_*p*_ ∪ *B*_*gt*_ is the union of these regions. Multiple detections are then ordered in terms of decreasing confidence. Multiple positive detections of the same region will only count the first detection as a positive and the rest as negative detections (False Negatives). If a ground-truth region contains no detections this counts as a FN also. There are no True Negative (TN) values as background regions are never actively identified. For a given class, a precision-recall curve is computed from a method ranked based on confidence. The precision is then calculated (precision = TP/(TP +FP)), and the recall (recall = TP/(TP+FN)) across all the data at each rank. We simplify the data by taking the maximum precision for any recall value, which results in the generation of a precision/recall curve, the area under which we can use to compare different algorithms. The so-called AP (Average Precision) metric is achieved my taking the maximum precision across all recall values and taking the average. Sometimes you will also see the metric mAP (mean Average Precision), this represents the mean AP value yielded from evaluation over different classes (i.e. performance across different cell types or objects).

Within the architecture of Faster-RCNN and YOLOv2 there are points at which randomness is injected into the training process. For example, the way the regions are selected and how they are shuffled for the training is done randomly. Much of this is unseeded, in that it is non-reproducible. This will mean every time a network is trained the resulting model will be slightly different, and the model will see different training data at different points of training. As a consequence the accuracy of trained models will vary slightly when retrained. If the variance is very high, it suggests that the training procedure is not well optimized for the problem. In this situation with relatively small amounts of data some variance is likely. To gain awareness of the network stability each experiment was repeated three times and the average precision measured, averaged and the standard deviation calculated.

### 3D acquisition algorithm

The AMCA (Automated Microscopy Control Algorithm) is written in python and is available in the following repository, https://doi.org/10.5281/zenodo.2594644).

Prior to the acquisition, the user defines the rough positions in which in the system scans for cells using the ‘collect_position.py’ script. With the script running, the user scans the slide manually and saves the positions of the stage at key-points around the area to be imaged. A minimum of four points is required to scan a rectangular area. At each location the user focuses the microscope on the cells using the z-piezo and stores the location using a python script. Once complete the algorithm interpolates the positions across the entire area, with a user defined sampling rate (e.g. every 200 μm). This array of spatial local forms the basis from which the acquisition of each stack takes place in the XY dimension.

For the ‘passive’ version of the algorithm, Micromanager is used to acquire stacks at each location defined using the ‘collection_positions.py’ script. Micromanager (v. 1.4.23) was configured to do this using the ‘PositionList’ functionality (a set of XYZ points) and was set to acquire stacks at these locations, for this work image stacks of 10 images sampled in ‘z’ at 0.5 μm intervals were acquired. The system then systematically acquires these stacks one-by-one and saves the output stacks to the computer as ‘tiff’ files. Using the ‘passive’ version of the AMCA algorithm, each stack is scanned and the location of the cells detected and exported using either the YOLOv2 or Faster-RCNN algorithms. The positions of the cells are stored in an output file and exported for subsequent processing.

Labview software (v. 15.0.1f10) was used with the ‘active’ form of the algorithm. In this mode, imaging and movement of the microscope act together dynamically. A python kernel running ‘amca.py’ was started before acquisition using the Python Integration Toolkit for Labview (Enthought). At regular intervals the Labview software would obtain an image from the microscope camera and then trigger the python function ‘analyzeAndMove’. The python function then analyses the image using either YOLO or Faster-RCNN, saves the image, and then if any cells are present will then trigger the microscope to move up in the ‘z’ dimension to the next position.

An image is then acquired in this location and Labview then activates the python function ‘analyzeAndMove’ again to analyze the image. If again cells are detected within the image, the image is saved and the microscope triggered to move up in ‘z’. This process is repeated until cells are no longer detected. At this point the microscope is instructed to return to the initial ‘z’ position in the stack and then to move down in the ‘z’ dimension until cells are no longer detected within this XY spatial location. At the point, the image stack is saved and the process repeated at the next XY location defined earlier.

Another option when using the ‘active’ mode is to visualize the cell localizations in real-time using the augmented reality system. For this modality the Labview software is set to use a python function called ‘analyzeAndView.py’ which allows the user to control the stage and visualizes the outputs the graphics for display using the augmented reality system. As the user moves the microscope specimen any cells detected will be shown surrounded by blue boxes.

An important aspect of both the ‘passive’ and ‘active’ forms of AMCA is its ability to extract the cells from volumes once detected in the individual slices; one cell detected in one z-slice, is not by default connected by reference to the same cell detected in the following slice. This connectivity problem is non-trivial to solve, as the detected cell regions do not necessarily overlap perfectly between the slices and may appear intermittently if the detection accuracy is low. To solve this challenge we modified a popular tracking algorithm called SORT (Simple Online and Real-time Tracker) [29] software and used it for linking the object detections between ‘z’ slices to form volumetric bounding regions for each detected cells. Algorithms such as SORT ensure that an object labeled in one image is connected to the same-labeled object in a subsequent frame. SORT is based on the principle of a Kalman filter, which means that it can accommodate significant perturbations to the cells. In both the ‘active’ and ‘passive’ AMCA variants SORT is applied offline, after acquisition but could easily be applied online also. Once cells bounding regions are uniquely identified through the volume they are saved as regions to the ‘tiff’ file metadata using a python library that encodes the regions in a format compatible with ImageJ/Fiji (for source, https://doi.org/10.5281/zenodo.2594733).

Subsequent analysis of images and regions acquired using AMCA were performed using ImageJ/Fiji and python scripts. All these scripts along with detailed instructions are available in the repository (https://doi.org/10.5281/zenodo.2594644).

## Results

### Initial Validation

Our goal for applying object detection networks to microscopy was ultimately so that these algorithms could be applied to find and isolate cells within a 3-D environment. As they stand, object detection algorithms are predominantly used to find objects in single 2-D photography images or in movies, and the training material is supplied to the algorithm exclusively in a 2-D format [9–15]. Single-plane images are far easier to label by users than 3-D volumes, requiring only a 2-D bounding box to be placed around examples within the image. Therefore, we wanted to establish our methodology for microscopy, including training and prediction, in 2-D and then apply it in a 3-D environment. To validate the object detection algorithms we created six different cell based datasets and modified the networks so that they could be trained on this data and also be validated against holdout test data (i.e. not used for training). Each dataset was divided into train and test datasets and the object detection networks were trained and evaluated on the data. With the exception of the neuroblastoma phalloidin data, each dataset was created and imaged within our host institution using conventional wide-field microscopes. The neuroblastoma phalloidin data was generated from an online resource [25] and the ground-truth segmentations converted into bounding box representations.

Figure 1 (and Figure S1) illustrates the six datasets generated for this study with ground-truth bounding boxes and predictions made by YOLOv2 and Faster-RCNN. First of all, we were interested to see if the technique would work at all. When applying vision based networks it is typical to inherit the weights of the convolutional layers from a pre-trained representation network, through a technique known as transfer learning [30]. The idea being that for visual tasks, many of the learnt features used for these applications are universal and will work with minimal tuning for a different objective. This holds true for identification of objects in images, but we were unsure whether the standard procedure of transfer learning would be sensible when starting with a network trained to identify objects in photography and then training it to recognize cells in microscopy images. In the case of Faster-RCNN the weights of the representation layers are taken from VGG16 [31] and for YOLOv2 they are taken from Darknet [32], both of which are pretrained on the photography image database ImageNet [33]. Upon visual inspection, the predictions made by YOLOv2 and Faster-RCNN on the test data were accurate with respect to the ground-truth regions created during the annotation of the datasets. This justified that the standard user protocol for applying object detection was applicable to application in microscopy without modification. Technically it is possible to train a network from scratch for this application, but for object detection this is not recommended due to the complexity of the overall network. Ideally a large corpus of data for cells would be generated and a representation network trained for classifying images into different cells types. This is not a small undertaking since the ImageNet database contains 14,197,122 images and 21,841 different categories. Fortunately however, it appears as though the VGG16 and Darknet pretrained networks form a good foundation on which to train the cell object detection capability. In the following result sections we quantified how accurate the predictions were and how they could be further optimized.

**Figure 1:**
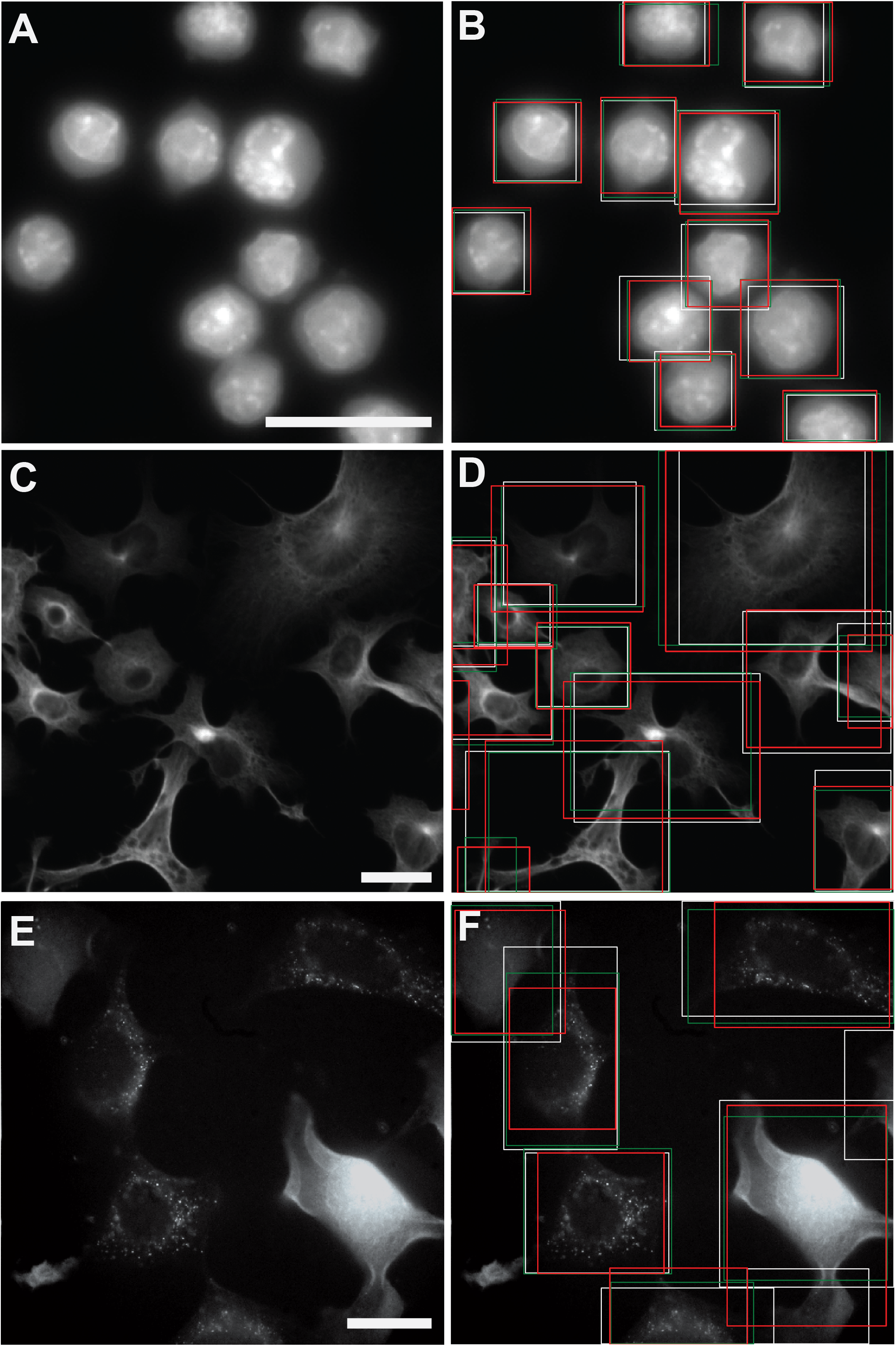
Example fluorescence microscopy data generated for our study with corresponding ground-truth human annotations and object detection predictions. **A-B**) eukaryotic cell dataset, stained with DAPI. **C-D**) neuroblastoma cells stained with GFP-phalloidin. **E-F**) HEK cells expressing GFP-SCP2 protein. Ground-Truth boxes, (**white**), YOLOv2 prediction boxes (**red**), Faster-RCNN prediction boxes (**green**). Scale bar (25 μm).

Machine learning algorithms that lack a closed-form solution generally require iterative training to get to an optimum solution that can be then used to make predictions. Neural networks are no exception to this: the weights of a neural network are adjusted and the performance evaluated many thousands of times before the optimum configuration of the network is found. Another potential issue with our initial validation was the concept of over-fitting. Over-fitting is a known and persistent problem within statistical methods, whereby the model being trained contains more parameters than can be justified by the amount and dimension of the training data. This problem is observable through assessment of the model on data unseen during training (i.e. test data), and this problem is often observed as a drop in accuracy past a certain point of training (i.e more than a certain number of iterations). With the six datasets (Figure 1) we evaluated the accuracy of the algorithm at different iterations of training. We found that typically, a peak level of accuracy was reached before the accuracy stabilized to a consistent value. For this study, we however found that overall, 20,000 iterations for Faster-RCNN (Figure 2) gave a stable and consistent high accuracy (i.e. the accuracy is high and does not fluctuate) across the test data for the basis of comparison, and for YOLOv2 (Figure S2A), 10,000 iterations a reliable stopping point to achieve the highest accuracy, and so all comparisons were made after this many iterations of training for each algorithm.

**Figure 2:**
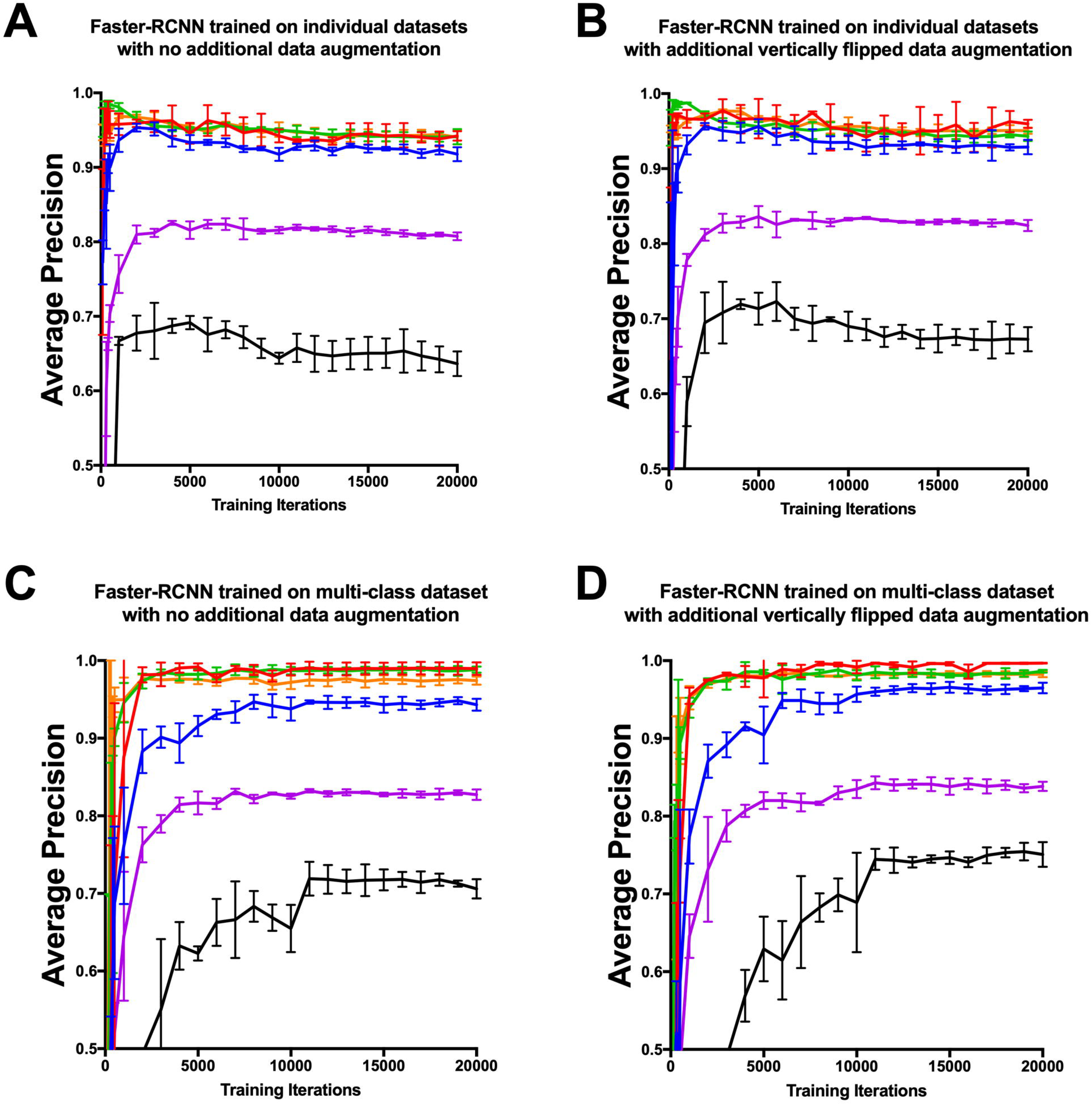
Average Precision at different levels of training for Faster-RCNN across six different datasets. **A-B**) Average Precision for Faster-RCNN network trained and evaluated on individual datasets without (**A**) and with (**B**) additional vertically flipped training data augmentation. **C-D**) Average Precision for Faster-RCNN trained across multiple datasets and evaluated on individual datasets without (**C**) and with (**D**) additional vertically flipped training data augmentation. Erythroblast DAPI cells (**blue**), neuroblastoma phalloidin dataset (**magenta**), fibroblast nucleopore dataset (**red**), eukaryote DAPI dataset (**orange**), C127 DAPI dataset (**green**) and HEK peroxisome dataset (black).

The first experiment performed was to train Faster-RCNN and evaluate its average precision (AP) on each of the six datasets. AP is a metric that evaluates the performance of the algorithm taking into account its accuracy and precision (see Materials and Methods for a full definition). This output performance is shown in Figure 2A at key training iterations, and the 10,000th iteration AP is summarized for each dataset in Figure 3A (cond. a). From this data it was clear that the algorithm performed better on certain datasets over others. This trend was seen throughout the study even with certain enhancements. The HEK peroxisome dataset was the most challenging (0.64 ± 0.02 AP ± SD), followed by the neuroblastoma phalloidin dataset (0.81 ± 0.01, AP ± SD), whereas Faster-RCNN performed very well on the fibroblast nucleopore, eukaryote DAPI and C127 DAPI datasets achieving >0.94 AP with no additional optimizations. For the same experiment YOLOv2 (Figure S2A and Figure 3B) performed better for the HEK peroxisome dataset (0.80 ± 0.02, AP ± SD) and the Neuroblastoma phalloidin dataset (0.89 ± 0.01, AP ± SD) compared to Faster-RCNN and >0.96 AP for the other datasets. Later, we definitively compare YOLOv2 and Faster-RCNN performance head-to-head with optimum settings for both algorithms. Interestingly the AP versus iteration number comparison (Figure 2A) showed that Faster-RCNN was clearly over-fitting the data. This is shown by the drop in performance in many of the datasets, with additional training iterations (i.e. the accuracy peaks and then declines with more iterations). This is not seen in the equivalent training profile for YOLOv2 (Figure S2A evaluated at the 10,000th iteration), indicating that over-fitting is not such a problem with this network and potentially explains some of the reason why YOLOv2 outperforms Faster-RCNN in general and on this data.

**Figure 3:**
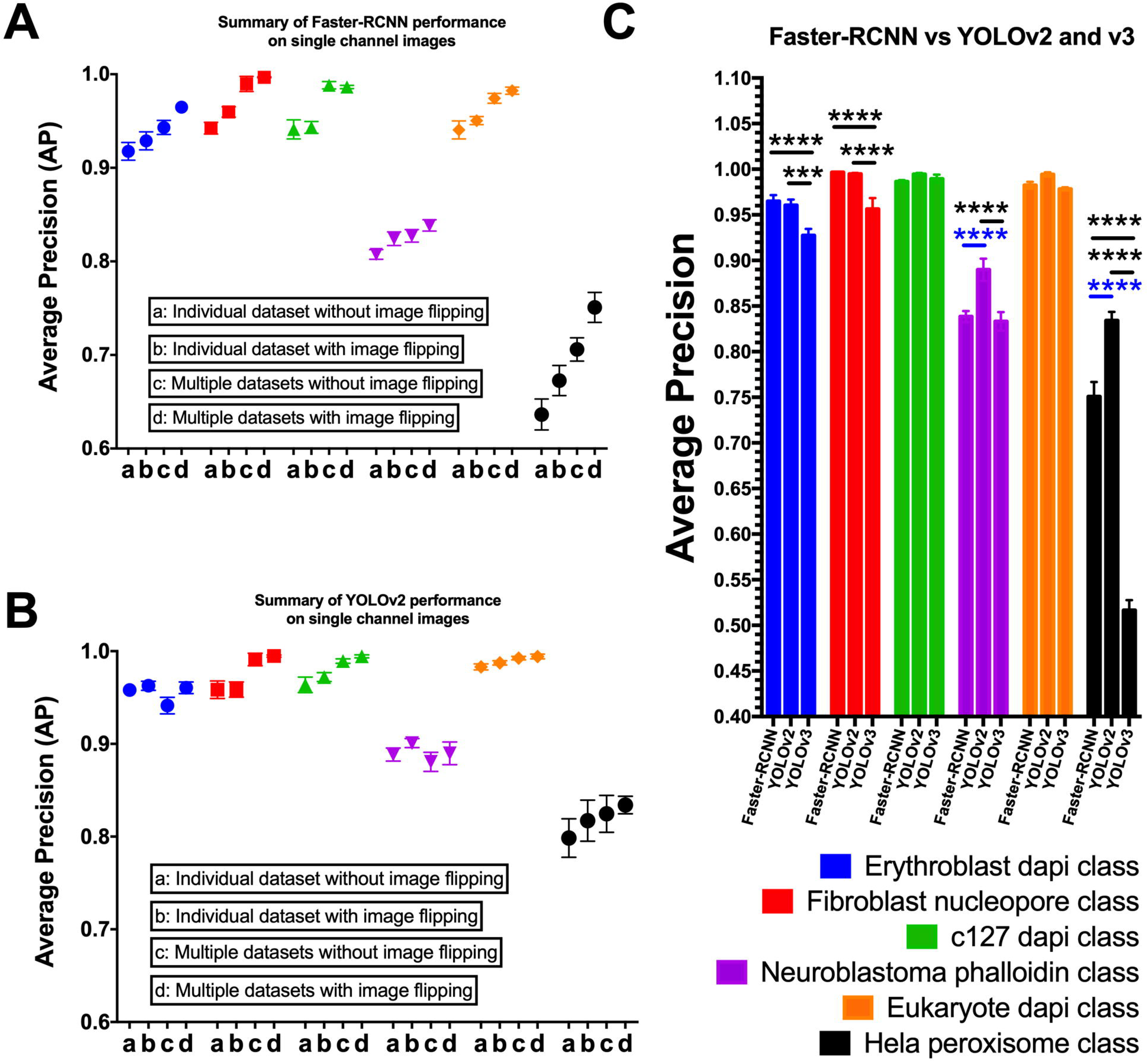
Summary comparison of object detection algorithms for cellular detection. Performance of Faster-RCNN (**A**) and YOLOv2 (**B**) when trained on: individual datasets without (**a**) and with additional flip data (**b**) augmentation and when trained using multiple datasets and without (**c**) and with (**d**) additional vertically flipped data augmentation **C**) Pairwise comparison of Faster-RCNN, YOLOv2 and YOLOv3 and trained on datasets with multiple classes and evaluated on single class with additional vertically flipped data augmentation. Erythroblast DAPI cells (**blue**), neuroblastoma phalloidin dataset (**magenta**), fibroblast nucleopore dataset (**red**), eukaryote DAPI dataset (**orange**), C127 DAPI dataset (**green**) and HEK peroxisome dataset (**black**). Statistical comparisons were made using 2-way ANOVA and Sidaks multiple comparison test and compared Faster-RCNN to YOLOv2 (blue) as well as comparisons between Faster-RCNN, YOLOv2 and YOLOv3 (black) for each dataset (n=3 in each case).

### Training across multiple classes

One of the key advantages of object detection algorithms is that they have been developed to classify objects across multiple categories and so are typically jointly trained to potentially recognize and differentiate a number of different objects simultaneously. Conventionally, in microscopy, we are only interested in localizing spatially a cell of particular type rather than classifying what type of cell it is, as the scientist who prepared the sample often knows this. Intuitively therefore, for microscopy data it would seem sensible to train networks on individual classes of data and then, as necessary, retrain the network from scratch to recognize subsequent datasets as needed. Instinctively, one would believe that this would create higher performing networks as the optimization is focused on boosting the accuracy for a particular class (cell type). As mentioned however, one of the challenges of machine learning is avoiding over-fitting, and this is clearly observed in the Faster-RCNN data as a decline in AP with iteration (Figure 2A). If we train our algorithm on several discrete, but similar datasets (in this case different cell classes) it is likely we can increase the amount of training data and produce an algorithm which generalizes better and performs more accurately on each individual cell class. We were interested therefore to know whether training across multiple classes of data simultaneously conferred an accuracy improvement over when the network was trained on only one specific dataset. Through training Faster-RCNN across all six datasets the AP increased on average 5.0% (0.90 mAP) (Figure 3A, cond. c) with respect to the equivalent model trained independently on each individual dataset (0.86 mAP, Figure 3A, cond. a). Interestingly, the over-fitting was no longer present when looking at the AP versus iteration profiles (Figure 2C, accuracy plateaus and does not decline). For YOLOv2 a smaller increase of 1.3% on average was seen in the AP for those models trained across multiple datasets (0.93 mAP, Figure 3B, cond. c) when compared to the precision for models trained on individual data (0.92 mAP) (Figure 3B, cond. a). In some cases for YOLOv2, a non-significant decrease was observed when training was performed across multiple datasets such as in the erythroblast DAPI and neuroblastoma phalloidin classes. It not clear why this may be, but it is clear that these datasets did not show much improvement under any condition, and so may represent a saturation in the performance of this type of network. In either case, training across multiple datasets boosts the performance of the framework being trained.

### Data augmentation

Object detection and many other computer vision algorithms that employ deep learning have some kind of data augmentation procedure to maximize the amount of data used for training. Data augmentation involves applying different kinds of transformations that alter the data subtly, providing informative variations of the training data. For example, rotation and scaling of objects provide clues as to how these objects may appear if positioned differently in test images, effectively increasing the number of relevant training examples for the model to learn from. This regularization method has been shown on numerous occasions to improve the accuracy of algorithms at test time. Object detection algorithms by default use data augmentation methods in order to increase the apparent number of images present.

Data augmentation is effective as long as the transformations are meaningful, for example if your dataset contained images of cars, it would not be meaningful for those images to be flipped vertically for example or the pixel intensities inverted. You are very unlikely to see a car upside down in photography and so a classifier doesn’t need to be trained on images simulating this. Flipping the image horizontally is fully permissible however, as you are just as likely to see cars coming from both sides. In this application the rules for augmentation are a little different as within microscopy it is fully permissible to flip the images horizontally and vertically and also to flip them both ways at the same time. This is because in conventional fluorescence microscopy the images are rotationally invariant (unlike in some other forms of microscopy e.g. DIC, phase contrast). In conventional fluorescence microscopy, images that are flipped vertically are just as realistic as those that are unflipped. Therefore, we were interested to see if we could improve the accuracy of the detection through flipping the images vertically to generate additional training data. The ‘flip’ conditions (Figure 3A and B (cond. b, d)) show the incremental improvement that augmenting the training data provides. In each dataset, for both Faster-RCNN and YOLOv2 when trained on a single dataset (Figures 3A and B, cond. b), or across multiple datasets (Figures 3A and B, cond. d) the additional flipping augmentation increased the accuracy of the resulting algorithm. The only exception to this was the C127 DAPI dataset with additional flipping (0.99 AP) which saw a marginal reduction when trained on multiple datasets and evaluated on the C127 dataset, compared to the unflipped data (0.98 AP). This difference was not significant. Therefore, given the bulk of the datasets and approaches it is clear that both Faster-RCNN and YOLOv2 benefit from additional vertical flipping of training data. This acts to increase the quantity of training data in a meaningful way improving the generalizability and accuracy of the resulting trained model.

### Faster-RCNN versus YOLOv2 and YOLOv3

From the described experiments it was possible to optimize the performance of both Faster-RCNN and YOLOv2. In both cases, the models trained on data that also included vertically flipped data and were trained across multiple datasets provided the best performance in each case. In terms of accuracy, Faster-RCNN and YOLOv2, once optimized, were very similar for most datasets (Fig. 3C). Later in the study YOLOv3 became available and so was included in the summary analysis [23]. Supplementary Table 1 summarizes the accuracy of either algorithm when evaluated on the datasets and Figure 3C summarizes the data graphically. In the more challenging HEK peroxisome and neuroblastoma phalloidin datasets the performance is significantly better with the YOLOv2 network compared to the Faster-RCNN algorithm (Figure 3C, blue statistical comparisons). In the rest of the data the results were indistinguishable and overall YOLOv2 was not significantly better than Faster-RCNN in terms of mAP. This shows that YOLOv2 is performing better on more challenging data, but with the correct optimizations both frameworks can perform near perfectly on less challenging data. The performance of YOLOv3 was disappointing on many of the datasets especially the more challenging ones and was performing significantly less well that the other two algorithms in all but the C127 DAPI and Eukaryote DAPI classes. The reason for this is not totally clear but is likely is due to the increased complexity of the network and likely requirements this puts on the level of training data required. At this stage, YOLOv2 is the better choice of algorithm with more challenging data and also has the added benefit of being a faster algorithm [21] than Faster-RCNN (67 FPS vs 7 FPS respectively) which makes it a more desirable choice.

### Multi-channel data

We were also interested in the possibility of using the object detection algorithm on multi-channel data. Both the neuroblastoma and the glycophorin A data were dual-stained and acquired within two different colour channels (Figure 4A and B). Each stain was discretely staining different aspects of the cell. In the case of the Neuroblastoma cells they were stained with phalloidin (green) for the actin cytoskeleton and DAPI (blue) for the cell nucleus, whereas in the case of the erythoblast cells they were stained for glycophorin A protein(green) and DAPI (blue) for the nucleus. We found that both YOLOv2 and Faster-RCNN were clearly capable of recognizing multiple channel images and the additional information was not detrimental to the classification performance. For the erythroblast two channel data (Figure 4C, cond. b), the performance was much the same as for the same images used with just the DAPI single channel staining (Figure 4C, cond. a). For the neuroblastoma dataset the performance of the algorithm when trained on the two channel data was improved in relation to the performance on the single channel data Figure 4C (cond. a and c vs. b and d; magenta data). This relationship persisted for models trained on only the neuroblastoma dataset (Figure 4C, cond. a and b) and also when trained using both neuroblastoma and erythroblast datasets combined (Figure 4C, cond. c and d). We also attempted to evaluate the models trained on multi-channel data on single-channel data. As expected, the algorithms performed poorly (data not-shown), as the model had a dependence on the additional colour channel, which was not present in this case. For example, an algorithm trained to recognize cells based both on phalloidin and DAPI staining will not perform well, if it sees phalloidin staining in place of the DAPI training. The algorithm cannot sensibly recognize in this context that the DAPI staining is in fact phalloidin and so its performance will suffer. This underwrites the necessity for models to be specifically trained for recognizing images with the same number of channels, ordered in the same sequence.

**Figure 4:**
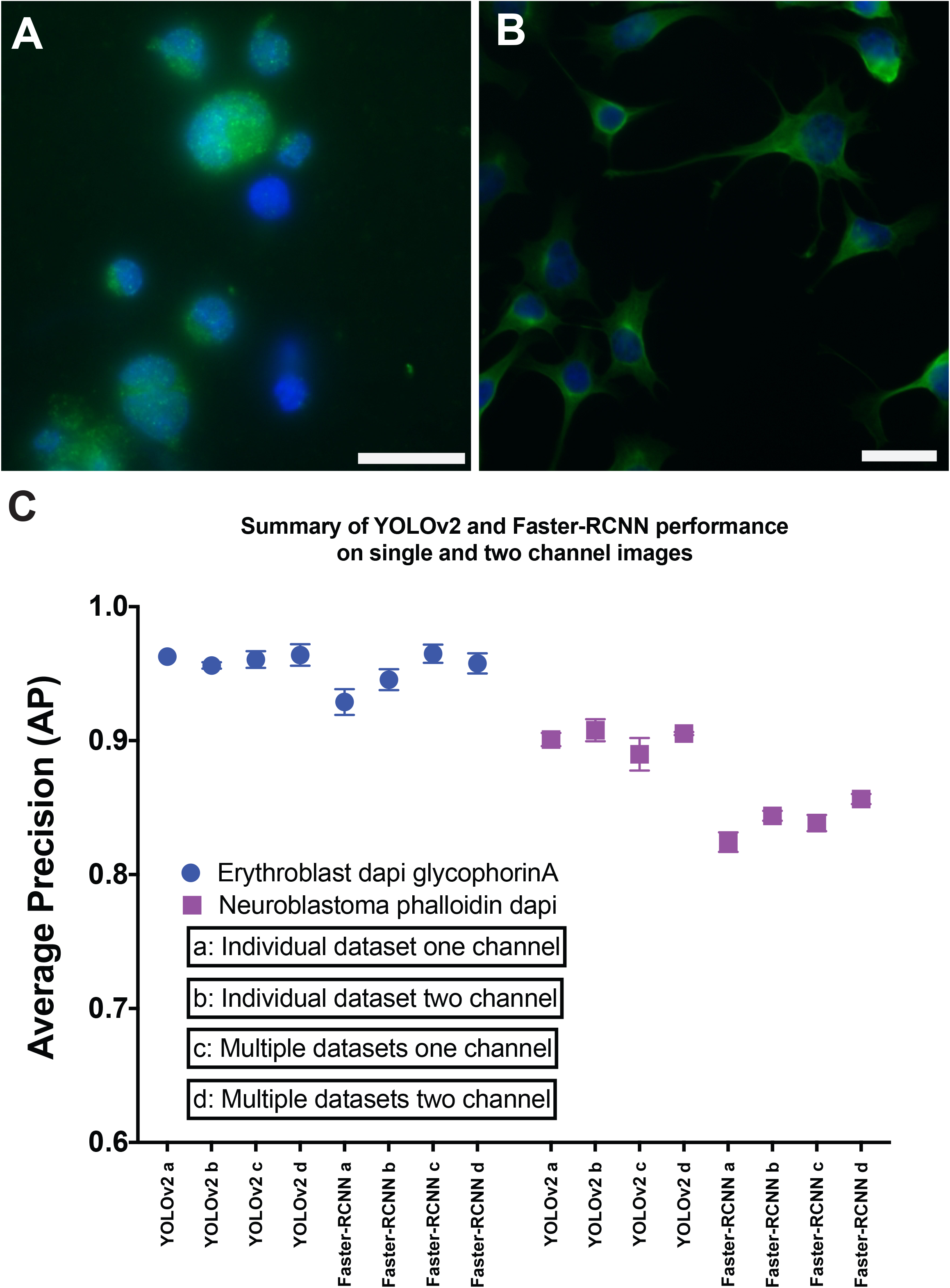
Summary performance of object detection algorithms when trained on multi channel data versus single channel data. **A**) Erythroblast cells stained with DAPI (**blue**) and for glycophorin A protein (**green**). **B**) Neuroblastoma cells stained with phalloidin (**green**) and DAPI (**blue**). Scale is 25 μm. **C**) Average precision of YOLOv2 and Faster-RCNN, comparing performance when trained and evaluated on a single dataset comprising of one channel data (**a**) and two channel (**b**) or when trained on multiple data comprising of one channel (**c**) and two chanels (**d**) respectively. All training data is additionally augmented through being vertically flipped. Erythroblast DAPI glycophorin A dataset (**blue**), neuroblastoma phalloidin DAPI dataset (**magenta**).

### 3D implementation of algorithm

Samples when viewed under a microscope are predominantly 3-D environments, where cells or specimens will often span more than one focal plane of the microscope. A natural progression from localizing cells in 2-D therefore, is to localize them in a 3-D environment. 3-D microscopy still predominantly involves taking 2-D images that optically section the 3-D volume of the sample. The image volume is then reconstructed subsequently and viewed by scrolling through one slice at a time, or through some form of rendering at an oblique angle. The challenge for cellular detection is to concatenate detections made in individual 2-D slices to create a 3-D region encompassing the entire cells. To adapt the object detection networks so that they could be used for acquiring cells in a 3-D environment, we developed the AMCA (Autonomous Microscope Control Algorithm). AMCA is a control framework which interfaces with microscope camera and control hardware to dynamically acquire images in 3-D. At its core is an object detection algorithm (either Faster-RCNN or YOLO), which informs the system whether there are cells present in a particular optical slice. AMCA comes in two forms, ‘passive’ and ‘active’ (See Figure 5 for graphical representation). The ‘passive’ algorithm, which is the simpler of the two algorithms, takes volumetric stacks of images which have been pre-acquired e.g. using Micro-manager software [34]. These stacks are then processed by the AMCA to find and label the sub-volumes occupied by the cells. This process is storage intensive and relatively slow as the images volumes are first acquired exhaustively across the slide before processing. The advantages of the ‘passive’ form however are its simplicity, because Micro-manager is easy to setup and run and freely available. The active form of AMCA works with Labview software [35] utilizing its depth of functionality. Through using custom scripts written in Labview and interfacing with python scripts it is possible to efficiently scan the slide and only acquire image volumes and slices where cells are identified. This is efficient, as only slices with cells in are retained and imaged and the microscope can move through areas without cells quickly and without continual prompt from the user.

**Figure 5:**
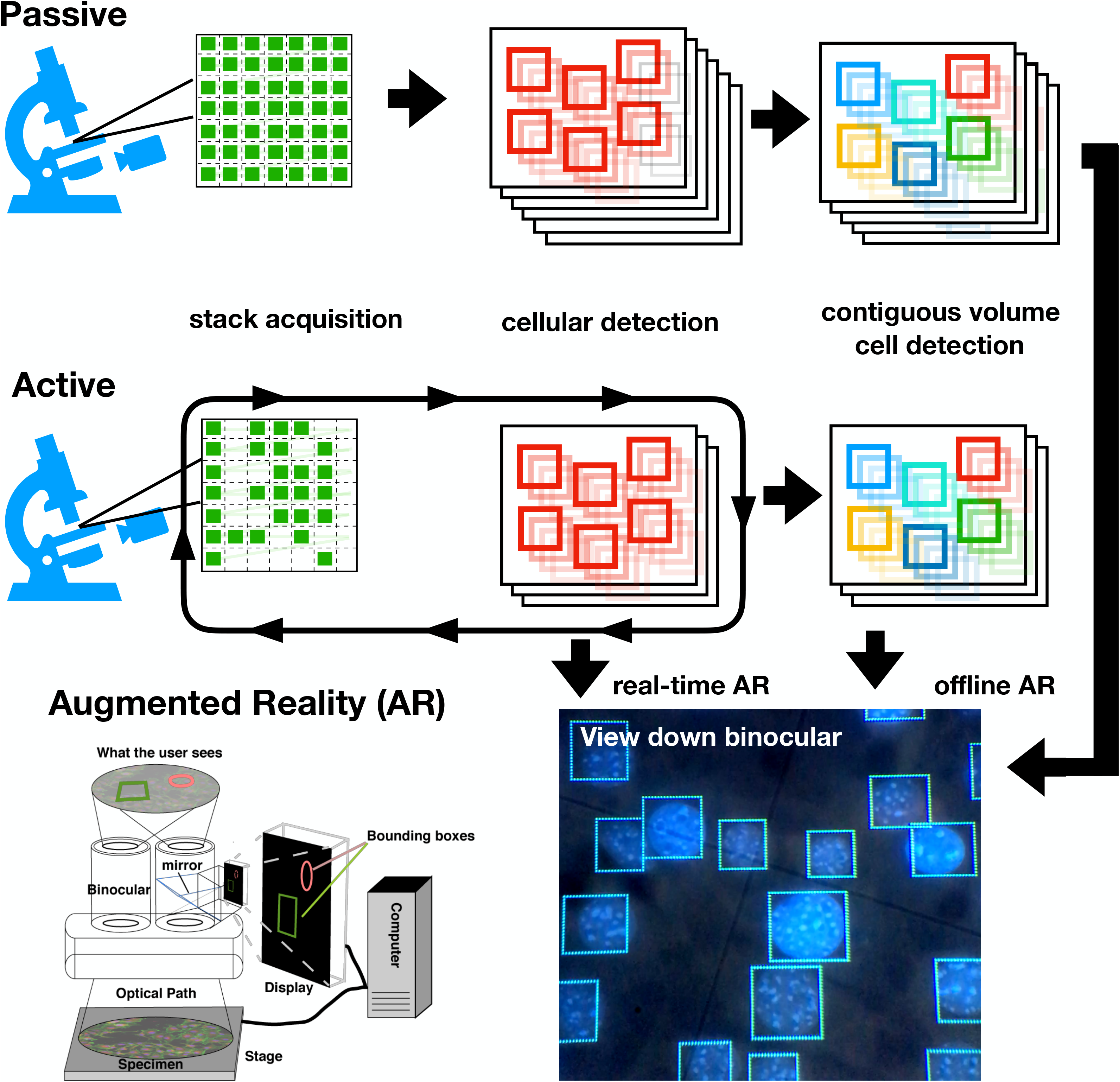
Schematic illustrating the differences between the ‘passive’ and ‘active’ forms of the AMCA algorithm. The ‘passive’ form of the algorithm involves exhaustively sampling the slide specimen and then analyzing the acquired stacks of images for the position of cells. Whereas the ‘active’ form of the algorithm involves imaging and analysis of positions in sequence. If a location contains cells, it is only scanned in ‘z’ dimension until no cells are detected, whereupon the microscope moves to the next position. In both modes, once all the regions containing cells have been saved they are analyzed to find the cells that are contiguous in the z-dimension, creating 3-D regions that encapsulate the cells. Augmented Reality (AR) allows the user to see the outputs of the analysis algorithm when viewing the sample down the microscope. The regions generated from cellular detection can be visualized through the augmented reality system in ‘real-time’, as the microscope acquires images, or subsequently ‘offline’, when the user views areas of the sample that have already been analyzed.

Although the AMCA system can be used to manually evaluate and acquire images it is most effective when set to automatically acquire images across the whole sample slide. As user input, it is necessary to specify a rough outline of the area to be inspected by the system. This process involves moving the stage to locations around the perimeter of the area to be imaged and selecting these locations to be bounding key-points. This prevents the system from going beyond the bounds of the slide and potentially damaging the system. The initial points to be imaged are then interpolated in x, y and z between these key-points. The regions can then be imaged densely or selected at random to sample the slide sparsely. When the system moves to a new location and an image is acquired in case cells are detected. In that case the microscope will move the sample up and then down in axial Z direction, acquiring 2-D images at each Z-position for 3D-reconstruction, and storing locations, until no cells are detected, at which point it signals to the microscope to move to the next spatial XY location.

Figure 6 shows the AMCA algorithm when applied to a volumetric acquisition of some C127 cells stained with DAPI. Figure 6A shows a single frame of the movie with the bounding boxes representing each unique cell localization and classification. The colour of each region shows the result of the tracking algorithm which discretely identifies each cell and does so for each classification made by the object detection algorithm. The classification persists through the in-focus region of the volume and each classification is linked with its counterparts from the same cell. Figure 6B shows each cell sub-volume detected and subsequently connected through ‘z’ by the AMCA tracking system (each unique colour shows a contiguous cell detection). These isolated stacks precisely represent the different z-slices of the cell across the image volume. Figure 6C shows a montage of the entire z-stack with the cellular detections depicted with discrete colours. The classifier has not been trained to recognize cells that are out-of-focus, and therefore out-of-focus frames are not retained. Our experiment clearly shows that object detection algorithms, when applied to microscopy, offer significant possibilities with regards to the identification and extraction of cellular sub-volumes.

**Figure 6:**
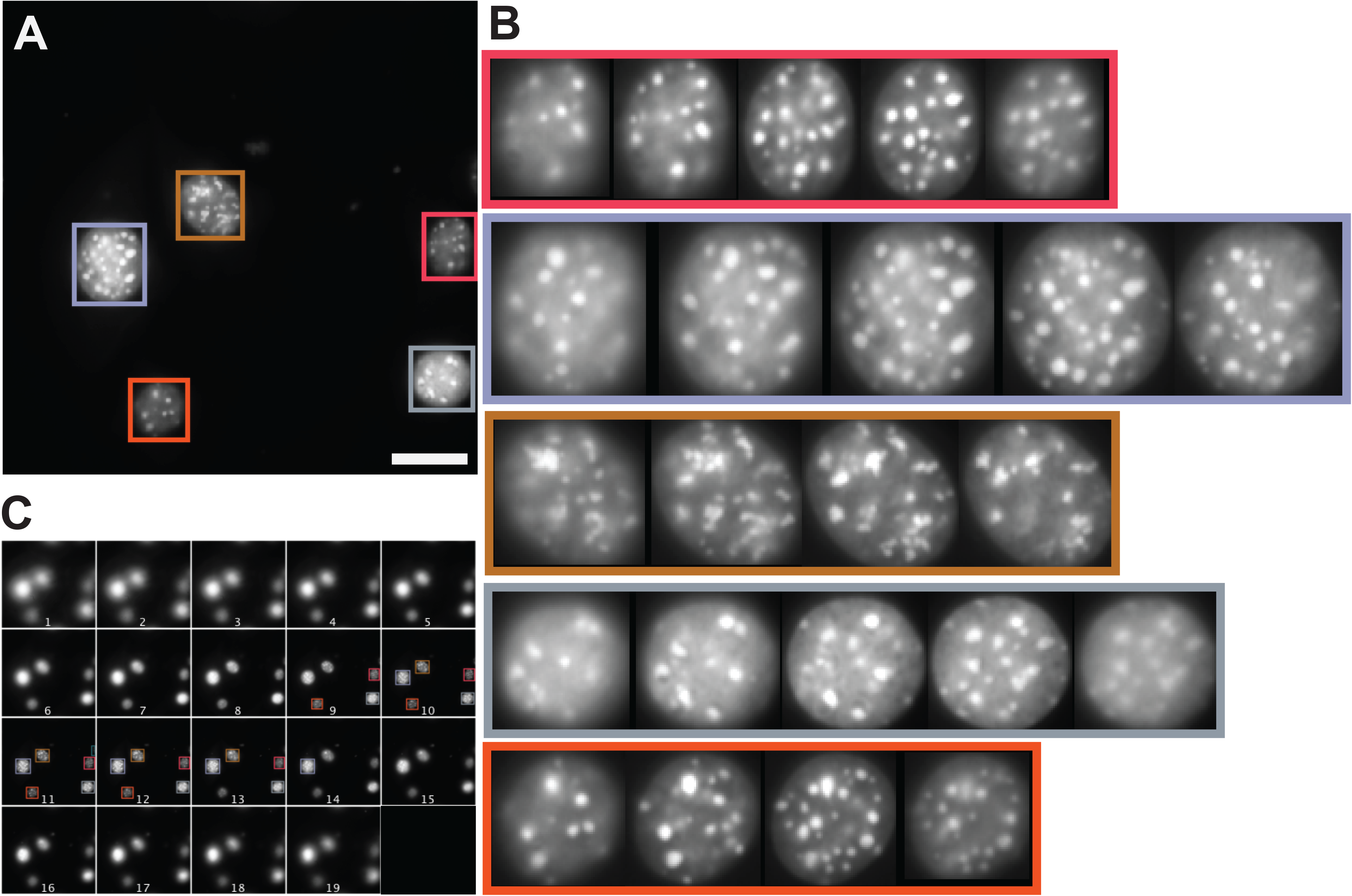
Acquisition of individual C127 cells from 3-D volumes. **A**) Example image with bounding boxes representing discrete cellular classification from Faster-RCNN and the colour represents track linking with adapted SORT algorithm. Scale is 25 μm. **B**) Depiction of cell classifications tracked through the different ‘z’ slices, colour border represents cells in ‘A’. **C**) Same cells as in ‘B’ but highlighted in original slices of full acquisition volume.

We wanted to showcase the potential for our method by performing a small-scale screen on cells and did so on a slide containing C127 cells, with nuclei stained with DAPI after having been treated with a technique known as RASER-FISH [26] (Figure 7). For this treatment cells are first incubated for 18 h with BrdU/BrdC. BrdU and BrdC are analogues for the nucleosides thymidine and cytosine and are incorporated into the DNA of proliferating cells. Almost all the cells should take up these analogues as they divide and synthesize new DNA. BrdU/BrdC are important for subsequent stages in the RASER-FISH technique, where they allow the generation of extensive regions of single-stranded DNA. A side-effect of RASER-FISH treatment is that conventional DAPI staining intensity is reduced in those cells that take up the BrdU/BrdC analogues. This is likely to be a twofold phenomenon; firstly DAPI fluoresces much more weakly when bound to single-stranded DNA compared to double-stranded DNA [36]). Further, DAPI preferentially binds adenosine-thymidine rich DNA regions, and therefore incorporation of BrdU, instead of thymidine, reduces the ability of DAPI to bind the cellular DNA and thus rendering the nucleus less fluorescent [37, 38]. For this study we made a screen encompassing 330 imaging positions spread across a slide area of (4.5×3.0 mm). Imaging positions were arranged uniformly and sparsely across the slide, 200 μm from each other. A low-resolution over-view image is shown in Fig 7A with a higher resolution image inset showing the imaged regions and cells identified in those regions (coloured boxes). In this case, any cells detected that touched the boundary were ignored. For the basis of overview statistics, image volumes were ‘maximum’ projected and the average intensity measured in each cell area. 2412 cells were detected and analyzed, and the procedure took 40 min in total. Figure 7B shows a summary histogram for the mean intensities of cells within the regions. The distribution has two peaks (33 and 55 intensity respectively) and suggests that there is more than one component contributing to the overall intensity distribution of the DAPI stained cells. Further experiments would be required to ascertain that this effect is real. Yet, our preliminary experiment shows the power of a higher-throughput automated approach to reveal effects not usually visible from assays run across a small number of cells. This data suggests the BrdU is not being taken up in every cell present, illustrated by the differential DAPI staining and resulting in a complex DAPI intensity distribution with multiple peaks. It is likely that those cells which appear to be brighter are those which have not taken up BrdU/BrdC during the incubation step. It would be interesting in subsequent work to establish whether the DAPI intensity in this manner can be used to ascertain the quality of RASER-FISH staining and whether this would be a good method for performing fully automated high-resolution imaging of specific cells.

**Figure 7:**
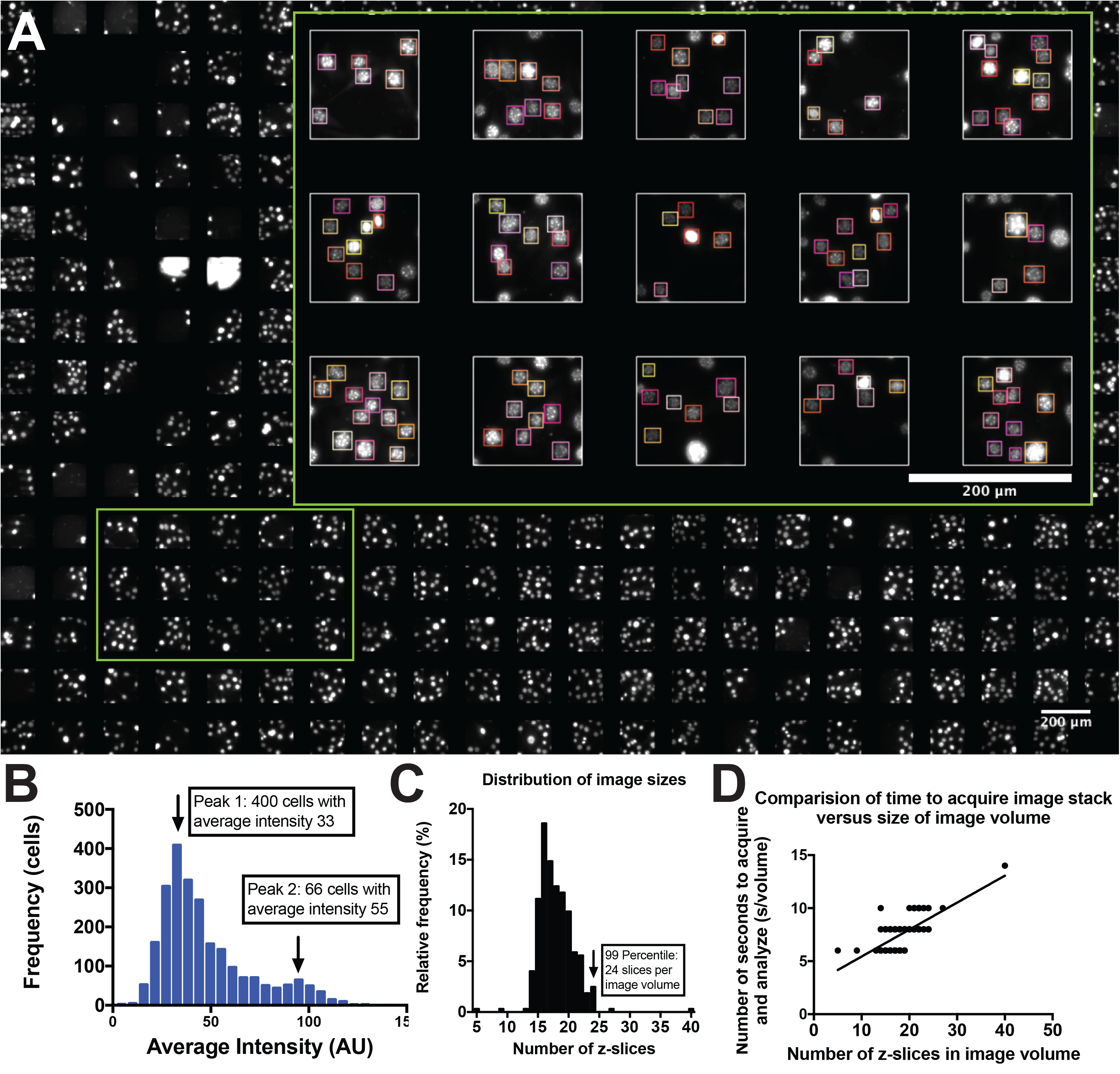
Preliminary screen of C127 cells stained with DAPI. **A**) low-resolution overview image of C127 cells acquired during screen. **Inset**) Zoom area shown by green rectangle. Image areas are shown with detected cells bounded by coloured boxes. Cells touching the image area boundaries are excluded in this analysis. **B**) Summary distribution for a histogram calculated over the mean intensity values for the cells acquired during screen. **C**) Distribution of image volume acquired size in terms of number of ‘z’ slices acquired during ‘active’ mode acquisition to encompass the cells present in each volume. **D**) Plot comparing the number of seconds required to acquire each volume at each position with the number of ‘z’ slices required for each volume. Straight-line represents regression line fit to data (slope:=0.4112, y-intercept=0).

Another interesting observation of the latter experiment is the time-taken to perform it. Rather than using the exhaustive ‘passive’ mode, the images were acquired and analyzed dynamically using AMCA in ‘active’ mode. Figure 7C shows a histogram summarizing the distribution of time-taken to acquire each image volume in seconds. This process is linear with respect to the number of slices present, as it takes longer to acquire and analyze image volumes that comprise more ‘z’ slices (Fig. 7D). On average it takes 7.5s to acquire a stack (of e.g. 18 slices) analyze the data, save the images, and move to the next position of the slide. Because the system is dynamic it cuts short the imaging of volumes where the imaging has already extended across all the cells present, saving valuable time. If we were to image exhaustively the same area, we would need image volumes comprising at least 24 frames (99 % percentile) to ensure we covered all the cells within the 3-D volume, assuming a symmetric positioning of cells centered on the start location. Image volumes comprising 24 ‘z’ slices would take at least 10 s per stack and 330 volumes would take 55 min in total compared to 40 min taken with the ‘active’ approach, which is almost a 1/3 quicker. For large screens this represents a large saving in time and also requires less storage. This screen was 6.09 GB in size compared to a 330×24 slice volumes which would be around 8.30 GB in size. This experiment proves that an automated system could easily localize and extract cells from a large area with very little human interaction and through doing it ‘actively’ one can save storage and acquisition time.

### Augmented reality acquisition

The augmented reality (AR) display gives real-time and off-line feedback to the user. The experience of using augmented reality in the context of microscopy enriches the experience and helps integrate the acquisition phase with the analysis of the sample. We demonstrate a simple solution to providing augmented reality using off-the-shelf components and optical elements (see methods for more details). The AR system lends itself very well to the fast analysis and processing of the tested object detection algorithms. The AR system can work in two different ways: ‘Real-time’ mode that works with the ‘active’ ACMA algorithm, where the system simultaneously analyses as you manually move around the slide. A user can quickly navigate around a slide and then receive analysis feedback regarding that area, identifying the cells and relaying information regarding brightness, number of foci, and size of cells. Whereas ‘Offline’ mode allows you to inspect the pre-acquired cellular positions on the slide with no-analysis over-head and so a slightly quicker frame-rate. The ‘offline’ analysis can be applied with data acquired through both the ‘passive’ and ‘active’ forms of the AMCA algorithm. Fig 5. shows a photo taken down the eyepiece that shows the augmented reality system working in ‘real-time’ mode. The image area is being imaged and relayed to the computer camera where the image is analysed and the regions are displayed and updated in real-time through the AR display adapter. For videos of the augmented reality working in real-time please refer to Supp. Mater. 3 and 4.

## Discussion

In this study we document the creation of a collection of datasets of cellular images and provide annotation in the form of bounding boxes. This has been an excellent resource for training, benchmarking and improving the object detection algorithms used in this study. This dataset is publically available and will be updated at regular intervals with additional data (https://zenodo.org/record/2548493). It is our hope it will become a valuable resource for future studies involving microscopy and object detection algorithms.

We have in this study thoroughly benchmarked two popular object detection algorithms, Faster-RCNN and YOLOv2, for the task of cellular classification and localisation. With the settings we have used we have found YOLOv2 to be a superior algorithm in terms of accuracy and speed. Additionally, we have found that both algorithms gave enhanced performance with additional data augmentation in the form of vertical flipping. We also found, as expected, that training across different classes simultaneously boosted performance of the networks, especially Faster-RCNN. We also benchmarked YOLOv3 toward the end of the study and surprisingly found the algorithm underperformed on multiple datasets. We believe this is due to the number of parameters and complexity that YOLOv3 has relative to YOLOv2 and how this inevitably requires more training material. YOLOv3 uses a more complex variant of darknet that forms the basis of the feature description layers. The YOLOv3 darknet has twice the number of convolutional layers (106 versus 53 for YOLOv2). A restriction of having more parameters is that you require more material to train adequately. Consequently, we believe that as networks get more and more complicated they will perform less well when applied on smaller-sized datasets, though interestingly in the case of our data, each of three algorithms perform well on the smallest dataset (C127 DAPI, 26 training images). The relationship is likely to be an interplay between the complexity of the data and the size of the datasets involved. It would suggest that unlike in the domain of photography where there is still a trend towards more complicated networks (and bigger and bigger datasets) that for microscopy we may see networks of a smaller size, which are easier to train. In either case the future for these kind of networks within the imaging sciences is likely to revolve around designing and generating network designs which can outperform YOLO and other networks, can function on smaller self-contained hardware, and can incorporate information such as focus into the classification process.

Bounding boxes, as a form of annotation, have the advantage that they are simple and quick to create and interpret. One criticism of using bounding boxes however is that cellular image analysis generally involves some form segmentation to discretize each cell, and to potentially reduce the impact of background pixels. Using bounding boxes however does not exclude this type of analysis and actually is an excellent prior for applying subsequent analysis methods through providing demarcation of the image and an appropriate region to provide as initialization. Conventionally, cells that are close to one another may upon segmentation appear connected. Traditionally a ‘watershed’ algorithm is used to separate the cells, but with the bounding box predictions it is possible to split the cells using the boundaries of the boxes as limits. Another use of bounding boxes is for Active Contours [39]. Active Contours are a popular form of statistical shape model whereby a parameterized spline is evolved through iteration, to fit a shape, which in this case would most likely be the perimeter of a cell. One of the main limitations of Active Contours and for many of the related level set or statistical shape models is the quality of initialization, especially where there are multiple objects present all which need to be fit. There is no generic way of initializing automatically despite this being critical to the successful application of this technique. Bounding boxes are an excellent method to bridge the problem of initialization and open up possibilities of application of these two techniques together. Furthermore, analyzing sparse foci distributions in scenes with dense cellular distributions of fluorescence can often be challenging as it challenging to separate the foci attributed to each cell. Using the bounding boxes however it is straightforward to delineate the cellular regions and attribute the foci accurately to each cell.

The AMCA algorithm shows that object detection algorithms have a valuable role to play in the development of automated microscopy acquisition algorithms. Although microscopes often operate within a 3-D environment, they most often only will acquire information as 2-D planes. As a consequence, 2-D object detection algorithms have a definite place in the acquisition process, with the possibility for 3-D classification and analysis post-acquisition. In this work, we have showed that AMCA can in combination with the pre-trained object detection algorithms, an automated stage and fast-acquisition camera can identify cells and acquire image volumes of those cells. This is a very powerful technique and easily trained by non-skilled users due to the simplicity of the training.

Following an acquisition session using the ACMA system the positions and dimensions of the cells are parameterized and stored. This data is rich and can be used in different ways. It would be possible, for example, to register the position of the slide during a series of imaging experiments. Often, the exact positioning and focus of a slide is lost when the slide is removed and then subsequently replaced on a microscope. This is often the case with conventional microscopy equipment and regular slide preparations. Due to the parameterization of the cells, it is possible with AMCA to accurately and quickly, register the slide during subsequent imaging. To do this a search strategy is employed which images a region of the slide and then matches the coordinate system of the two slides, before imaging the whole slide in the corrected coordinate system. Such developments would make it very easy to perform time-series experiments or to return to samples after an initial imaging experiment for repeated or different measurements. Work of this nature empowers users of conventional microscopes to perform large-scale acquisition and analysis experiments in the domain of conventional microscopy. This means that conventional slide preparation and regular microscopy equipment can be used to perform high throughput experimentation. We expect to utilise this technique for a number of applied experiments in the future, where the automation and recognition capability will boost the number of samples acquired and enable new types of experiments to be performed.

It is not essential to use an automated stage to yield the benefits of AMCA. Cells under a microscope can still be classified and tracked using the AMCA algorithm even with the stage under human control. The drawback is that when acquiring image volumes, there will not be a record of the z-plane position. As a consequence it works well as means of highlighting cells but not for recording 3-D volumes of cells. This functionality as is, however, works excellently as an augmented reality microscope system, annotating samples, as the user navigates around the sample looking for cells of certain brightness for example.

AR in the microscopy domain has several advantages over conventional visualisation of data: E.g. computer-based exploration of 3-D data can often be cumbersome and unintuitive. In contrast, a microscope is a well-designed tool for navigating a 3-D space and lends itself well to 3-D exploration. By augmenting the visual output of the microscope, the data is displayed in the context of the slide and so is enjoyable to view and explore in this way. It can sometimes be easier for a user to appreciate a sample when looking down a microscope. The user can also quickly review areas that haven’t been imaged with those that have. By having information overlaid on the image data one can understand it better.

## Funding

We would like to acknowledge the UKRI BBSRC (BB/P026354/1) and the UKRI MRC (MR/S005382/1a, MC_UU_12009) for support of this project as well as the Deutsche Forschungsgemeinschaft (Research unit 1905 “Structure and function of the peroxisomal translocon”), the EPA Cephalosporin Fund, the MRC (grant number MC_UU_12010/unit programs G0902418 and MC_UU_12025), the Wellcome Trust (grant 104924/14/Z/14 and Strategic Award 091911 (Micron)), MRC/BBSRC/EPSRC (grant MR/K01577X/1), the Wolfson Foundation (for initial funding of the Wolfson Imaging Centre Oxford), and the John Fell Fund.

## Acknowledgements

We thank John Prentice (Workshop Manager, University of Oxford) for his skilled technical and engineering input. We would also like to thank Silvia Galiani and Iztok Urbancic (HIU, University of Oxford) for their advice whilst setting up the microscope within the Nano-Immunology lab, as well as the Wolfson Imaging Centre Oxford and Christoffer Lagerholm for general support.

