## Supplementary Materials 1 for "Object Detection Networks and Augmented Reality for Cellular Detection in Fluorescence Microscopy Acquisition and Analysis"

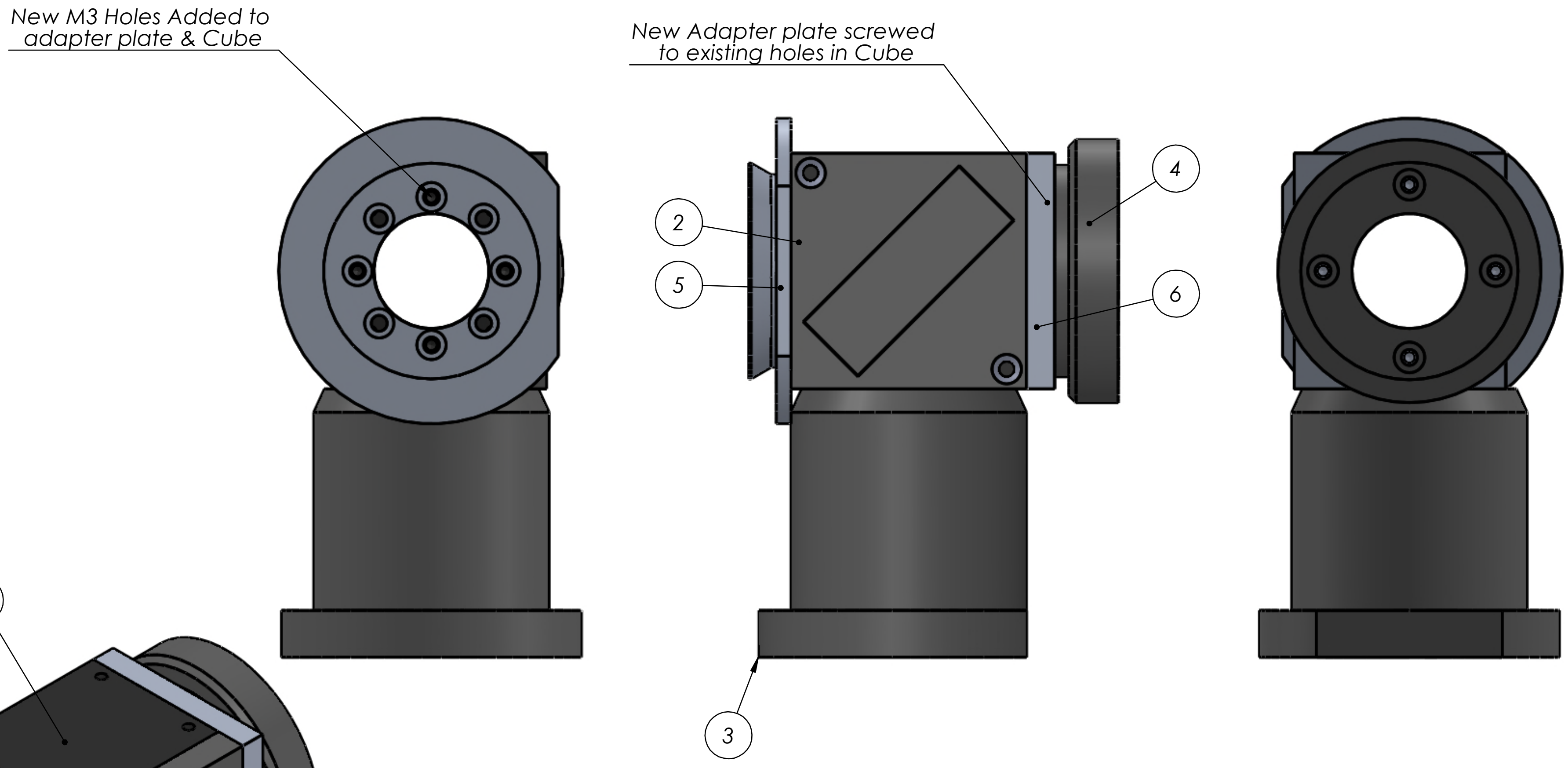


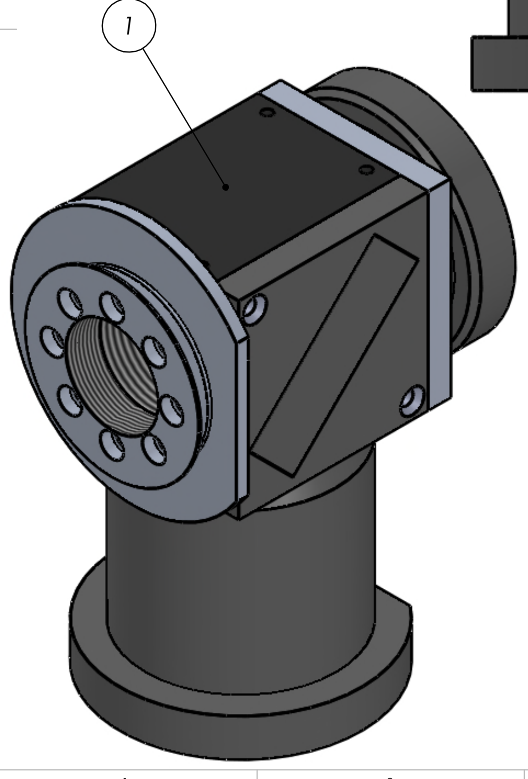


| ITEM NO. | PART NUMBER | DESCRIPTION | QTY. |
|---|---|---|---|
| 1 | DSI-CUBE-OL-UA_Cube | WIMM_ Computational Biology _Modified Part | 1 |
| 2 | DSI-CUBE-OL-UA_Cube_Plate | | 1 |
| 3 | DSI-CUBE-OL-UA_Cube_Cylindrical Clamp Post | | 1 |
| 4 | DSI-CUBE-OL-UA_Cube_Adapter Ring A | | 1 |
| 5 | DSI-CUBE-OL-UA_Cube_Adapter Ring B | Modified part | 1 |
| 6 | DSI-CUBE-OL-UA_Cube_Adapter Plate 1 | Manufactured part | 1 |

UNLESS OTHERWISE SPECIFIED:
DIMENSIONS ARE IN MILLIMETERS
SURFACE FINISH:
TOLERANCES: ISO 2768 class f
LINEAR: 0.5 up to 6 ±0.05
over 6 up to 30 ±0.1
over 30 up to 120 ±0.15
over 120 up to 400 ±0.2
ANGULAR: 1 deg

FINISH:

DEBUR AND BREAK SHARP EDGES

DO NOT SCALE DRAWING

REVISION A-01

DESCRIPTION:

| | NAME | SIGNATURE | DATE |
|---|---|---|---|
| DESIGN | jprentice | | 25/10/2018 |
| DRAWN | | | |
| APPV'D | | | |
| MFG | | | |
| Q.A | | | |
| Vendor | | | |
| Vendor Number | | | |

TITLE:

### MIGHTEX_Cube Adaptor

MATERIAL:

DWG NO.

A3

WEIGHT:

SCALE:1:2

SHEET 1 OF 1

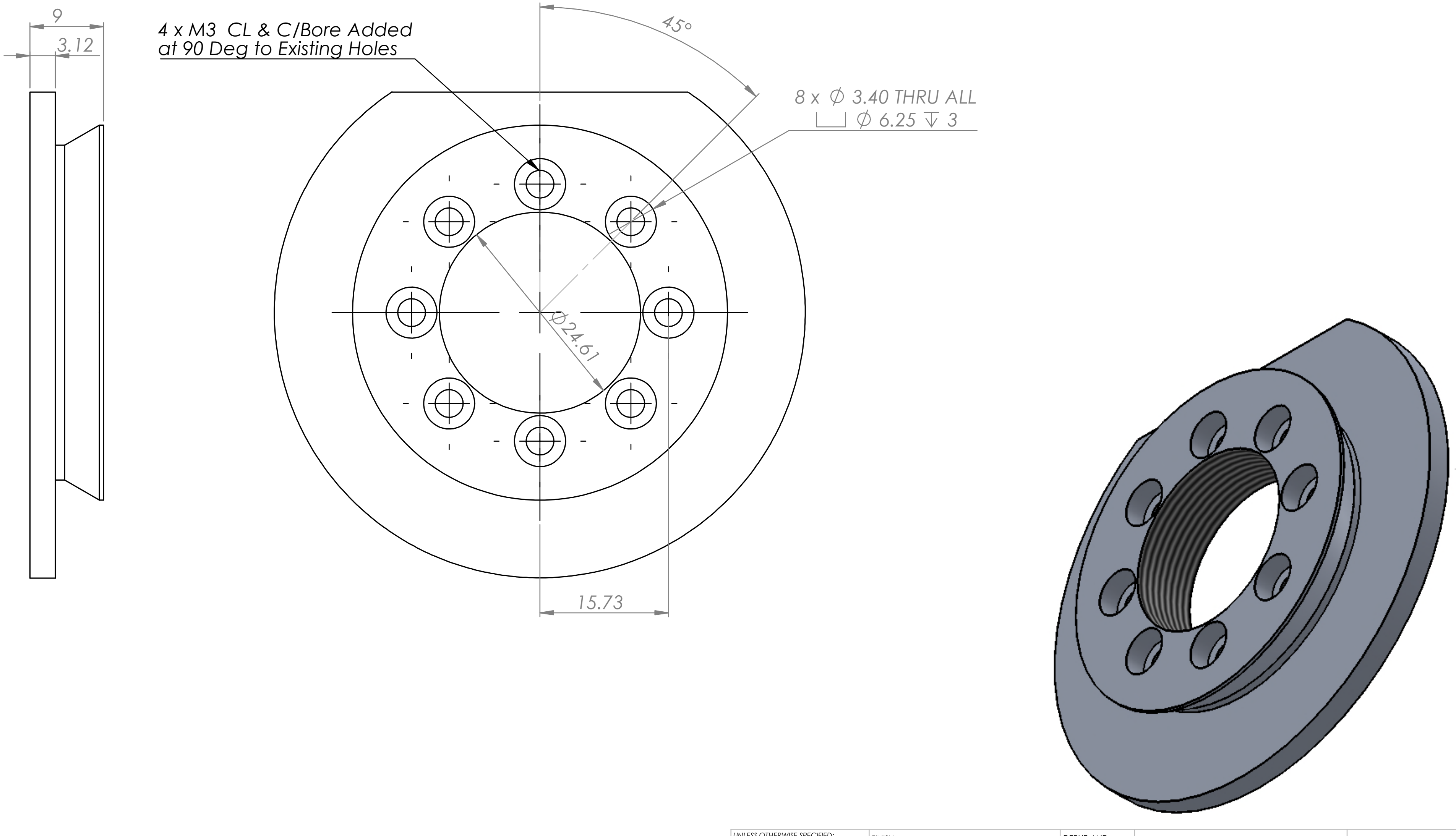


| | | | | | | | |
|---|---|---|---|---|---|---|---|
| UNLESS OTHERWISE SPECIFIED:<br>DIMENSIONS ARE IN MILLIMETERS<br>SURFACE FINISH:<br>TOLERANCES: ISO 2768 class f<br>LINEAR: 0.5 up to 6 ±0.05<br>over 6 up to 30 ±0.1<br>over 30 up to 120 ±0.15<br>over 120 up to 400 ±0.2<br>ANGULAR: 1 deg | FINISH: | | | | | DEBUR AND BREAK SHARP EDGES | DO NOT SCALE DRAWING — REVISION A-01<br>DESCRIPTION: Modified part |
| | NAME | SIGNATURE | DATE | | | | TITLE: DSI-CUBE-OL-UA_Cube_Adapter Ring B |
| DESIGN | | | 24/10/2018 | | | | |
| DRAWN | | | | | | | |
| APPV'D | | | | | | | |
| MFG | | | | | | | |
| Q.A | | | | MATERIAL: 1060 Alloy | | | DWG NO. 000-2111 — A3 |
| Vendor | | | | | | | |
| Vendor Number | | | | WEIGHT: | | | SCALE:1:1 — SHEET 1 OF 1 |

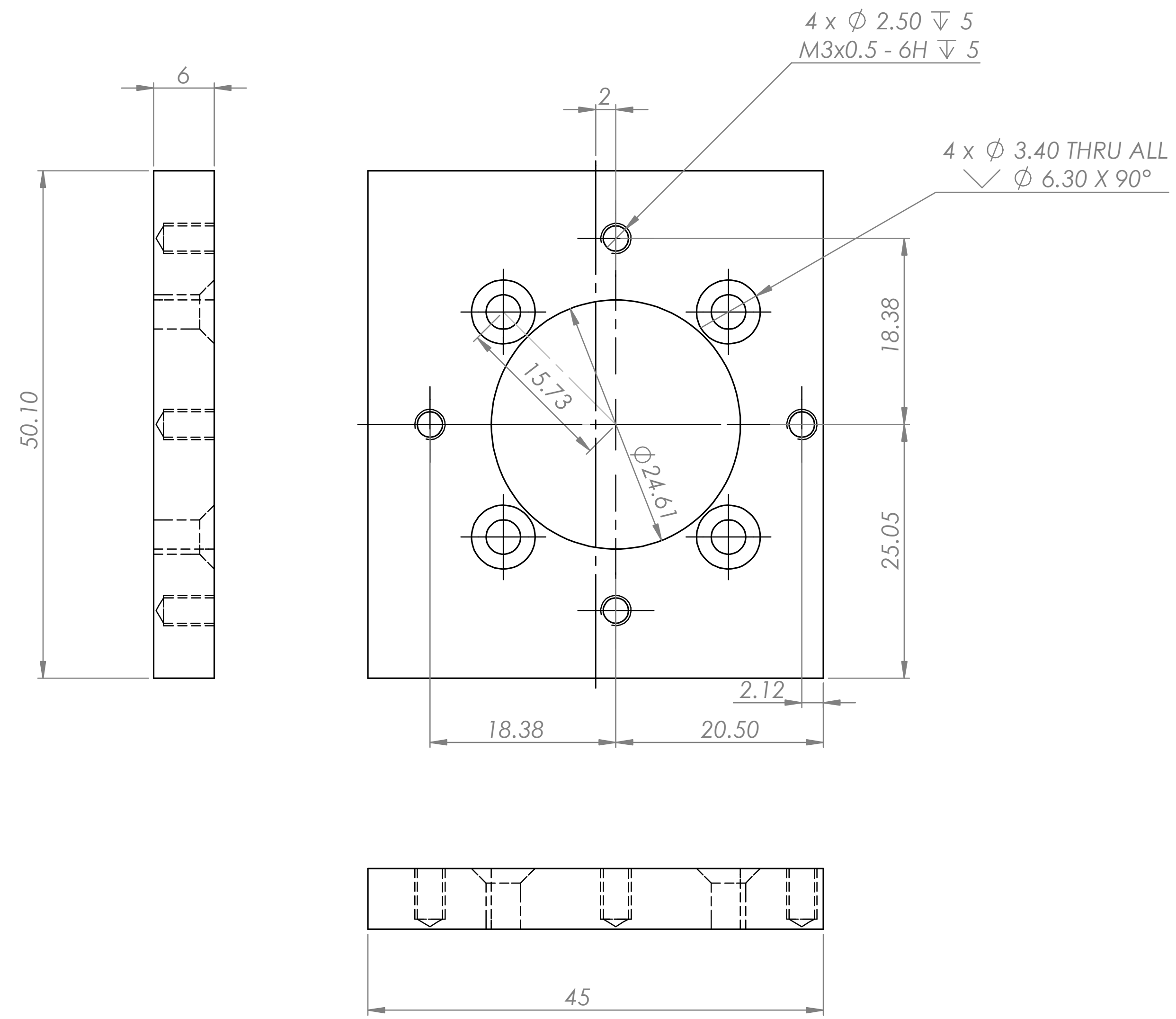


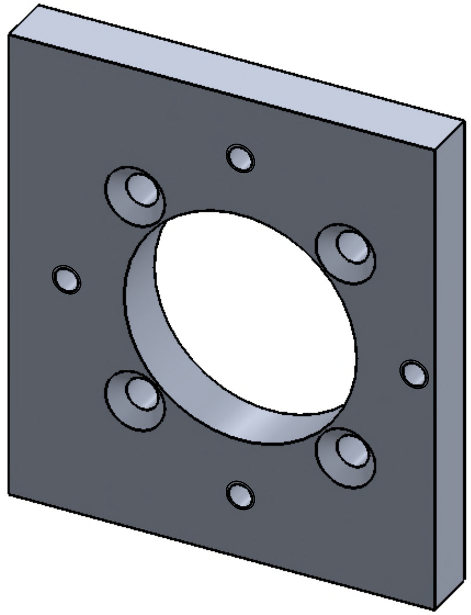

UNLESS OTHERWISE SPECIFIED:
DIMENSIONS ARE IN MILLIMETERS
SURFACE FINISH:
TOLERANCES: ISO 2768 class f
LINEAR: 0.5 up to 6 ±0.05
over 6 up to 30 ±0.1
over 30 up to 120 ±0.15
over 120 up to 400 ±0.2
ANGULAR: 1 deg

FINISH:

DEBUR AND BREAK SHARP EDGES

| | NAME | SIGNATURE | DATE |
|---|---|---|---|
| DESIGN | | | 25/10/2018 |
| DRAWN | | | |
| APPV'D | | | |
| MFG | | | |
| Q.A | | | |
| Vendor | | | |
| Vendor Number | | | |

DO NOT SCALE DRAWING

REVISION A-01

DESCRIPTION: Manufactured part

TITLE: DSI-CUBE-OL-UA_Cube_Adapter Plate 1

MATERIAL: 1060 Alloy

DWG NO. 000-2112

A3

WEIGHT:

SCALE:2:1

SHEET 1 OF 1

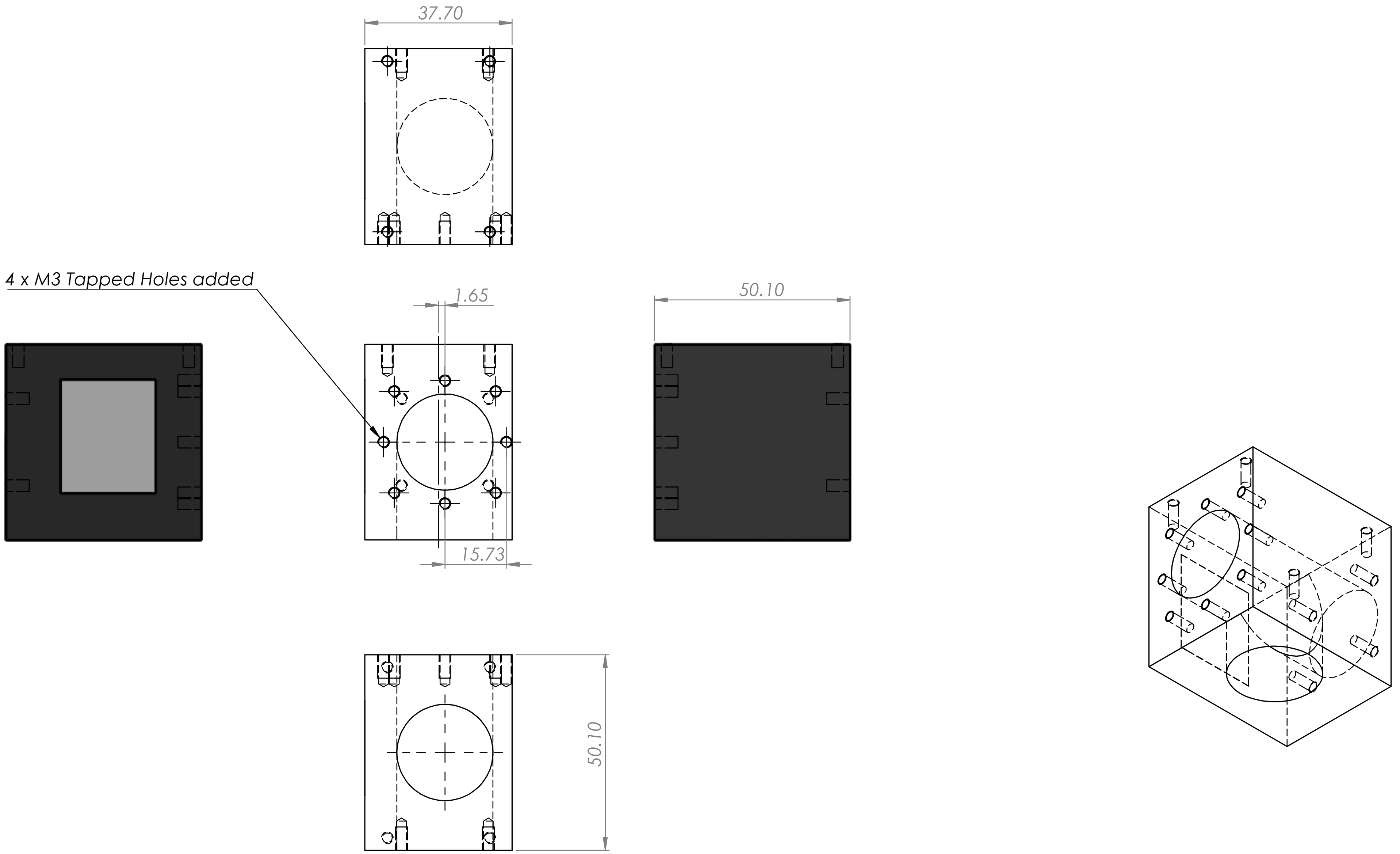


UNLESS OTHERWISE SPECIFIED:
DIMENSIONS ARE IN MILLIMETERS
SURFACE FINISH:
TOLERANCES: ISO 2768 class f
LINEAR: 0.5 up to 6 ±0.05
over 6 up to 30 ±0.1
over 30 up to 120 ±0.15
over 120 up to 400 ±0.2
ANGULAR: 1 deg

FINISH:

DEBUR AND BREAK SHARP EDGES

DO NOT SCALE DRAWING

REVISION A-01

DESCRIPTION:
WIMM_ Computational Biology _Modified Part

| | NAME | SIGNATURE | DATE |
|---|---|---|---|
| DESIGN | | | 25/10/2018 |
| DRAWN | | | |
| APPV'D | | | |
| MFG | | | |
| Q.A | | | |
| Vendor | | | |
| Vendor Number | | | |

TITLE:
DSI-CUBE-OL-UA_Cube

MATERIAL:
1060 Alloy

WEIGHT:

DWG NO.
000-2113

A3

SCALE:1:1

SHEET 1 OF 1
