## Supplementary Materials 2 for "Object Detection Networks and Augmented Reality for Cellular Detection in Fluorescence Microscopy Acquisition and Analysis"

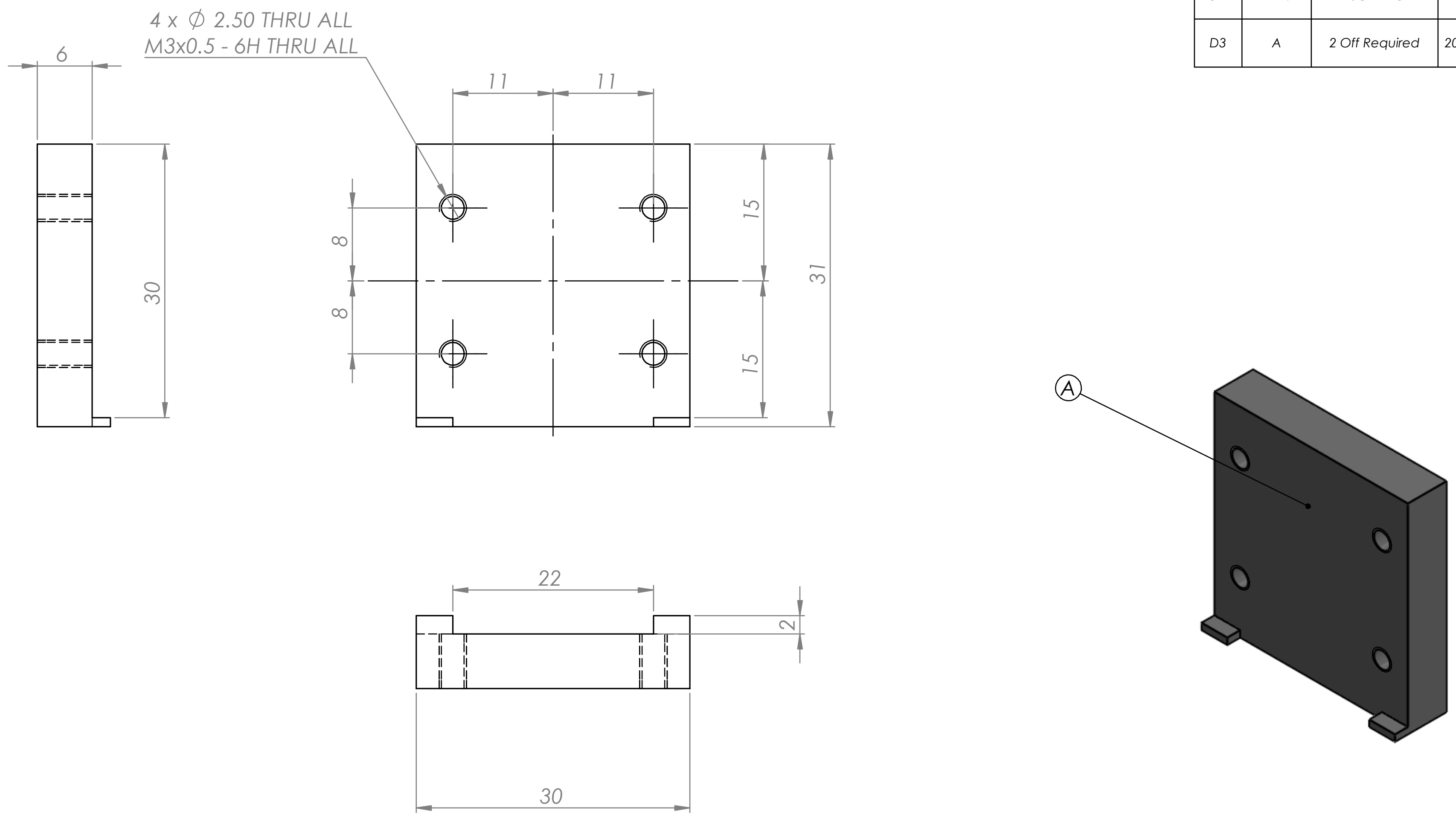

| REVISIONS | | | | |
|---|---|---|---|---|
| ZONE | REV. | DESCRIPTION | DATE | APPROVED |
| D3 | A | 2 Off Required | 20/11/2018 | |

UNLESS OTHERWISE SPECIFIED:
DIMENSIONS ARE IN MILLIMETERS
SURFACE FINISH:
TOLERANCES: ISO 2768 class f
LINEAR: 0.5 up to 6 ±0.05
over 6 up to 30 ±0.1
over 30 up to 120 ±0.15
over 120 up to 400 ±0.2
ANGULAR: 1 deg

FINISH:

DEBUR AND BREAK SHARP EDGES

DO NOT SCALE DRAWING

REVISION A-02

DESCRIPTION:

| | NAME | SIGNATURE | DATE |
|---|---|---|---|
| DESIGN | jprentice | | 19/11/2018 |
| DRAWN | | | |
| APPV'D | | | |
| MFG | | | |
| Q.A | | | |
| Vendor | | | |
| Vendor Number | | | |

MATERIAL: 40% GF PPS

WEIGHT:

TITLE: WIMM_30 x 30 x 30 Prism_Adapter Plate

DWG NO. 000-2176

A3

SCALE:2:1

SHEET 1 OF 1
