## Supplementary Figures for "Object Detection Networks and Augmented Reality for Cellular Detection in Fluorescence Microscopy Acquisition and Analysis"

| | Faster-RCNN | | YOLOv2 | | YOLOv3 | |
|---|---|---|---|---|---|---|
| | AP | SD (AP) | AP | SD (AP) | AP | SD (AP) |
| Erythroblast DAPI | 0.965 | 0.007 | 0.961 | 0.006 | 0.927 | 0.007 |
| Fibroblast nucleopore | 0.997 | 0.001 | 0.995 | 0.001 | 0.956 | 0.012 |
| C127 DAPI | 0.986 | 0.002 | 0.994 | 0.002 | 0.990 | 0.005 |
| Neuroblastoma phalloidin | 0.838 | 0.006 | 0.890 | 0.012 | 0.833 | 0.010 |
| Eukaryote DAPI | 0.982 | 0.004 | 0.994 | 0.003 | 0.978 | 0.002 |
| HEK peroxisome | 0.751 | 0.016 | 0.834 | 0.009 | 0.517 | 0.011 |
| **Mean AP (mAP)** | **0.920** | **0.006** | **0.945** | **0.006** | **0.867** | **0.008** |

Supplementary Table 1: Summary performance of Faster-RCNN and YOLOv2 and YOLOv3 algorithms on test datasets. AP, Average Precision, Number of samples =3, SD refers to standard deviation.

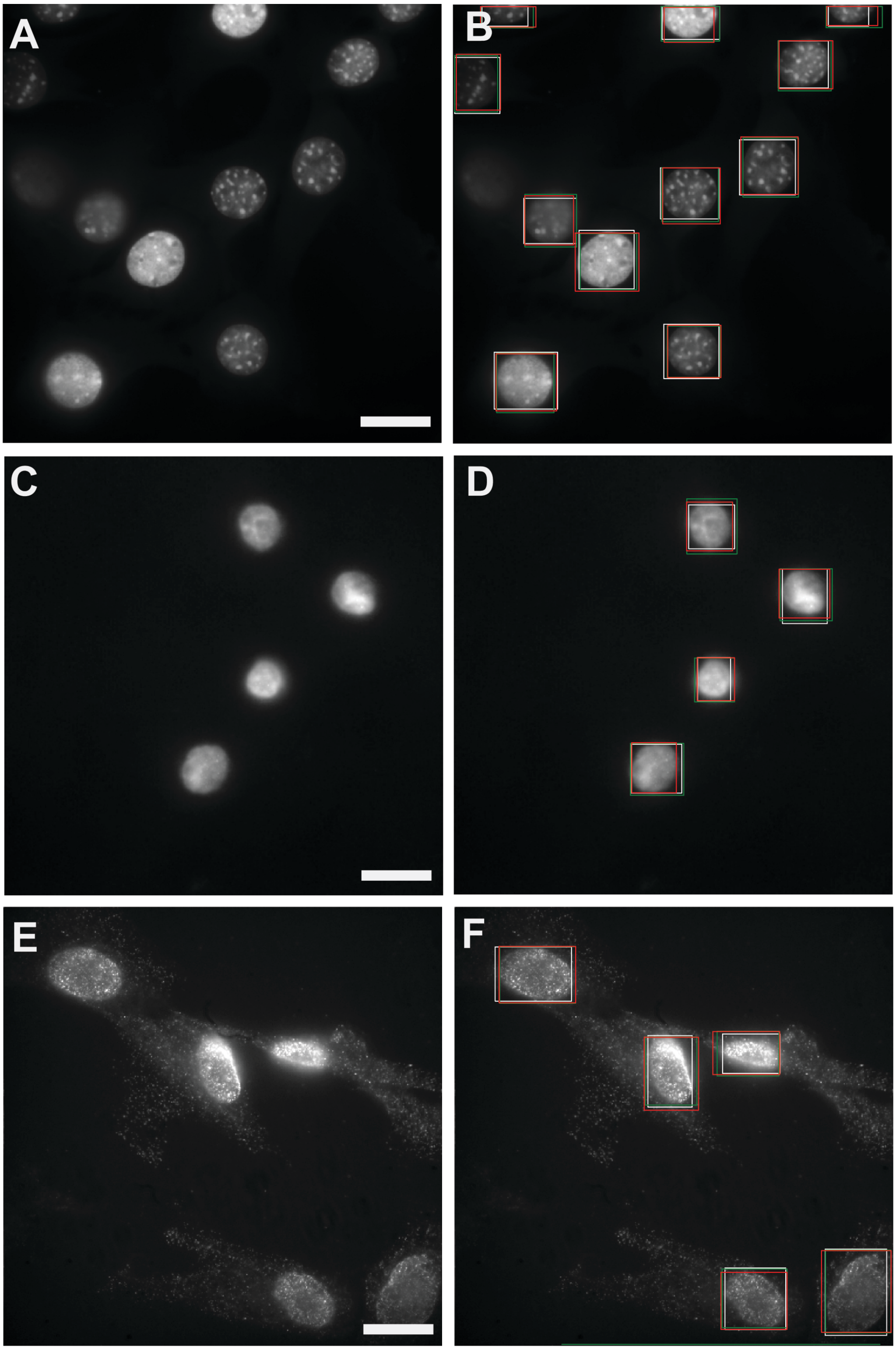


Figure S1: Example data generated for study with corresponding ground-truth human annotations and object detection predictions. **A-B**) C127 cell dataset, stained with DAPI. **C-D**) Erythroblast cells stained with DAPI. **E-F**) Fibroblast cells stained for a nucleopore protein. Ground-Truth boxes, (white), YOLOv2 prediction boxes (red), Faster-RCNN prediction boxes (green). Scale bar (25 μm).

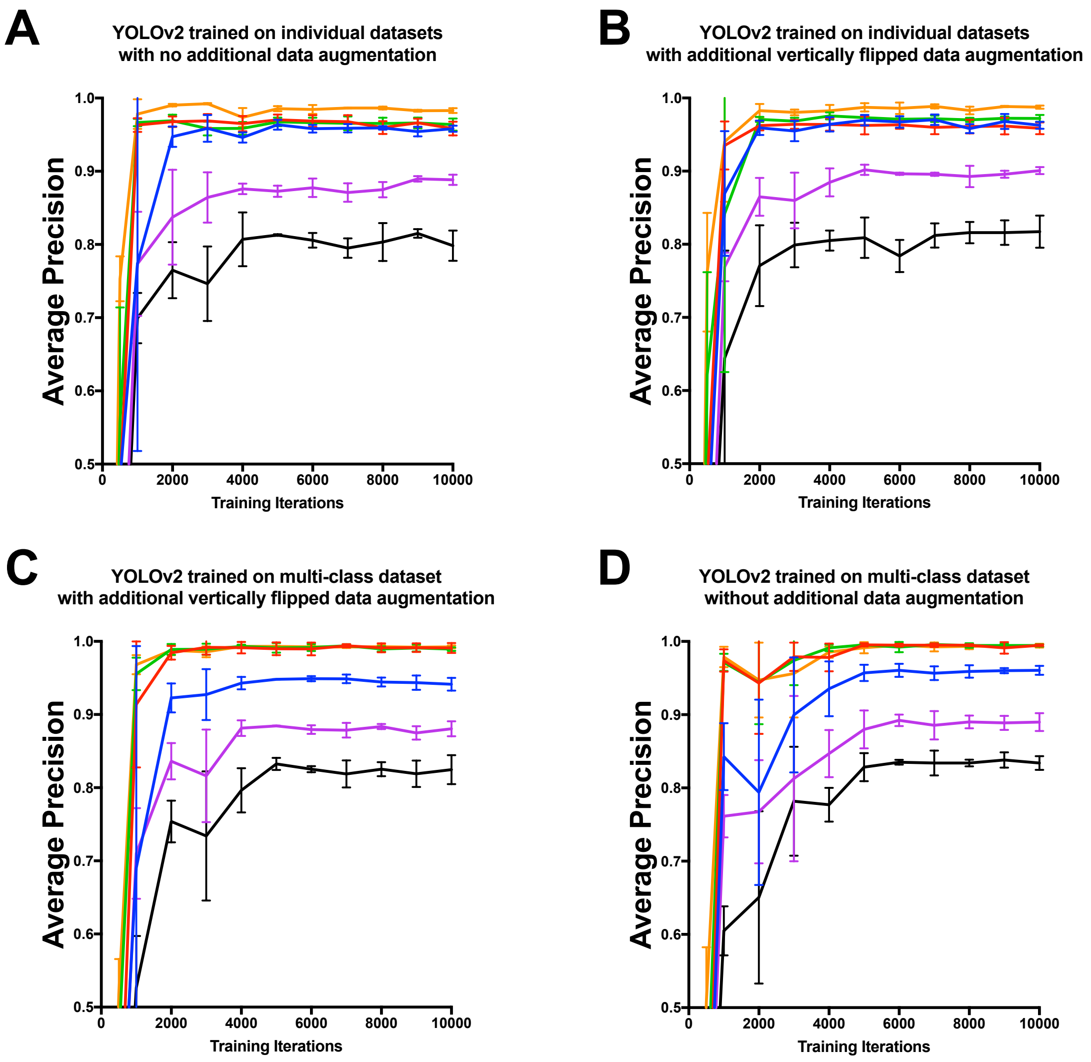


Figure S2: Average Precision at different levels of training for YOLOv2 across six different datasets. **A-B**) Average Precision for YOLOv2 network trained and evaluated on individual datasets without (**A**) and with (**B**) additional vertically flipped training data augmentation. **C-D**) Average Precision for YOLOv2 trained across multiple datasets and evaluated on individual datasets without (**C**) and with (**D**) additional vertically flipped training data augmentation.
